# Replicable generation of rhesus macaque iPSCs for *in vitro* modeling of genetic frontotemporal dementia

**DOI:** 10.64898/2026.03.17.712482

**Authors:** Julia C. Colwell, John P. Maufort, Kathryn M. Williams, Allison T. Makulec, Melissa V. Fiorentino, Jeanette M. Metzger, Heather A. Simmons, Puja Basu, Kerri B. Malicki, Celeste M. Karch, Jacob A. Marsh, Marina E. Emborg, Jenna K. Schmidt

## Abstract

At the Wisconsin National Primate Research Center, we have identified a family of rhesus carrying the *microtubule-associated protein tau* (*MAPT*) R406W mutation linked to frontotemporal dementia (FTD). Rhesus induced pluripotent stem cells (RhiPSCs) derived from these monkeys present a unique opportunity for *in vitro* modeling and comparison with cells derived from *MAPT* R406W human carriers. Here, we report the development of a reproducible method to generate RhiPSCs compliant with the standards of the International Society for Stem Cell Research (ISSCR) to support *in vitro* modeling of FTD*-MAPT* R406W. Our stepwise approach identified efficient methods for fibroblast derivation, fibroblast reprogramming to RhiPSC, and RhiPSC maintenance over continued culture. To derive fibroblasts from *MAPT* wild type (WT) and R406W monkeys, a combination of manual processing and overnight enzymatic digestion was required to maximize the number of low passage fibroblasts available for reprogramming. Fibroblast reprogramming to RhiPSC using Sendai viral vectors versus oriP/EBNA1 episomal plasmids revealed the latter as most efficient. Electroporation conditions for oriP/EBNA1 reprogramming were optimized to maximize plasmid uptake and cell survival. Ultimately, eight RhiPSC lines were derived from 4 donor rhesus monkeys (n=2 WT, n=2 R406W; two clonal lines per donor) and fully characterized according to ISSCR standards. RhiPSC stemness and genetic stability was best maintained on mouse embryonic fibroblast feeders in Universal Primate Pluripotency Stem Cell medium, as opposed to Essential 12 medium supplemented with IWR1, which produced cytogenetic abnormalities. Rhesus neural progenitor cells were generated using a monolayer protocol and expressed PAX6 and NESTIN after 21 days of differentiation. Our reliable method will be useful to labs seeking to derive RhiPSCs for preclinical studies. Overall, the RhiPSCs generated from *MAPT* R406W carriers will be a critical resource for evaluating the molecular underpinnings of tau-related neurodegeneration across primate species.

## 1.0 Introduction

Frontotemporal dementia (FTD) is the most common form of dementia in patients under 60 years of age (Bang et al., 2015). Mutations in the *microtubule associated protein tau* (*MAPT)* gene have been linked to FTD, including the autosomal dominant mutation *MAPT* R406W. Patients with *MAPT* R406W dementia clinically present with progressive Alzheimer’s disease-like cognitive impairment and personality changes (Reed et al., 1997). Furthermore, R406W tau filamentous inclusions are structurally similar to the paired helical filaments found in Alzheimer’s disease (Qi et al., 2025). Human induced pluripotent stem cells (hiPSC) generated from *MAPT* R406W carriers are valuable for analyzing tau-related FTD pathogenesis, understanding the genotype-phenotype relationship *in vitro*, and testing target engagement of therapies. Studies in hiPSC-derived neurons from these patients found that the *MAPT* R406W mutation is associated with increased tau cleavage, accumulation of misfolded tau, microtubule destabilization, disruption of mitochondrial transport, and mitochondrial dysfunction (Imamura et al., 2016; Mahali et al., 2022; Minaya et al., 2023; Nakamura et al., 2019). Moreover, bulk RNA sequencing of *MAPT* R406W hiPSC-derived neural progenitor cells identified alterations in molecular pathways involved in calcium homeostasis and synaptic function (Jiang et al., 2018; Minaya et al., 2023).

At the Wisconsin National Primate Research Center (WNPRC), we have identified a family of rhesus macaques carrying the same *MAPT* R406W mutation found in human FTD. These monkeys, currently being deeply phenotyped (results to be published elsewhere), present a unique opportunity to derive rhesus iPSCs (RhiPSCs) for *in vitro* studies of genetic FTD across species. Importantly, the RhiPSCs can be used for testing potential therapies before *in vivo* preclinical studies in these valuable monkeys. However, methods for RhiPSC derivation and maintenance are not as efficient when compared to those established for human iPSCs (hiPSCs), which can be produced from patients of any age or genetic condition (Karch et al., 2019). RhiPSCs were first derived in 2008 from rhesus macaque fibroblasts via retrovirus-mediated introduction of Yamanaka factors OCT3/4, SOX2, KLF4, and c-MYC (Liu et al., 2008). Several research teams have since reported the successful generation of RhiPSCs, yet publications vary in their reprogramming methods and conditions to maintain the resulting RhiPSCs over continued culture. Furthermore, few studies have derived more than one RhiPSC line (Hallett et al., 2015; Tao et al., 2021), highlighting the need for a robust and reproducible method for the derivation and maintenance of monkey iPSCs. Lastly, while publications of newly generated hiPSCs are expected to fulfill the reporting standards of the International Society for Stem Cell Research (ISSCR; (Ludwig et al., 2023)), studies of monkey iPSCs typically lack information such as cryopreservation methods or cell line authentication, underscoring the need to uphold similar standards across species to ensure reproducibility.

Here, we established a reproducible method of rhesus fibroblast reprogramming to RhiPSC using non-integrating oriP/EBNA1 episomal plasmids that resulted in eight RhiPSC lines derived from four animals (two clonal lines per animal), including two animals carrying *MAPT* R406W. In this report, we included both successful and unsuccessful reprogramming trials to illuminate the range of variables considered at each stage of our stepwise approach to develop a replicable method of RhiPSC generation. The RhiPSCs were characterized according to ISSCR standards following at least one freeze-thaw cycle, demonstrating cell stability and thus a robust maintenance method. RhiPSCs (one clonal line per animal) were patterned to rhesus neural progenitor cells (RhNPCs) to further demonstrate the differentiation potential and enable future studies on mechanisms of tau-related neuropathology in genetic FTD.

## 2.0 Materials and Methods

### 2.1 Ethics Statement and Overall Experimental Design

This study was performed in strict accordance with the recommendations in the Guide for the Care and Use of Laboratory Animals of the NIH (8th edition, 2011) in an Association for Assessment and Accreditation of Laboratory Animal Care International (AAALAC)-accredited facility (WNPRC, University of Wisconsin-Madison). The experimental protocols were approved by the Institutional Animal Use and Care Committee at UW-Madison (protocols #G006226-R01-A05 and #M005991-R02-A04). All efforts were made to minimize the number of animals used and to ameliorate any distress. An overall workflow of the study is depicted in **Figure 1**. All media formulations are provided in **Supplementary Table 1**.

**Figure 1.**
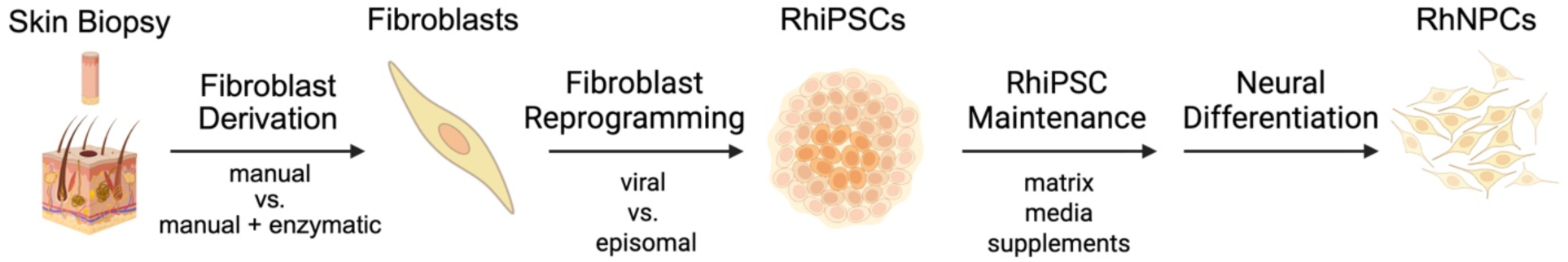
Overall experimental workflow with variables tested at each step. Abbreviations: RhiPSC, rhesus induced pluripotent stem cells; RhNPC, rhesus neural progenitor cells. Created in https://BioRender.com.

### 2.2 Tissue and Blood Collection

Skin biopsies were obtained from *MAPT* R406W (n=6 biopsies; 2 males, 2 females, 1.1-18.8 years) and *MAPT* WT (n=5 biopsies; 1 male, 2 females, 1.4-19.0 years) rhesus macaques (**Table 1**). For the procedure, the animals were anesthetized with ketamine (5 mg/kg IM) and dexmedetomidine (0.015 mg/kg IM) and their lower abdominal or upper back region was shaved. The skin was first cleaned with iodine, followed by isopropyl alcohol, and then flushed with sterile saline. Two pieces of tissue, each 6 mm in diameter, were excised using an Integra Miltex disposable biopsy punch (Integra, cat. no. 33-36) from the lower abdomen or upper back. Skin biopsies were immediately submerged into biopsy medium warmed to 37°C prior to transfer to the laboratory for processing. Biopsy sites were closed with 4-0 vicryl reverse cutting suture, and then lidocaine (0.05 mL/site SQ) was injected at the skin biopsy sites. Meloxicam (0.2 mg/kg SQ) was administered post-biopsy collection for pain and inflammation. Anesthesia was reversed with atipamezole (0.15 mg/kg IM).

**Table 1:**
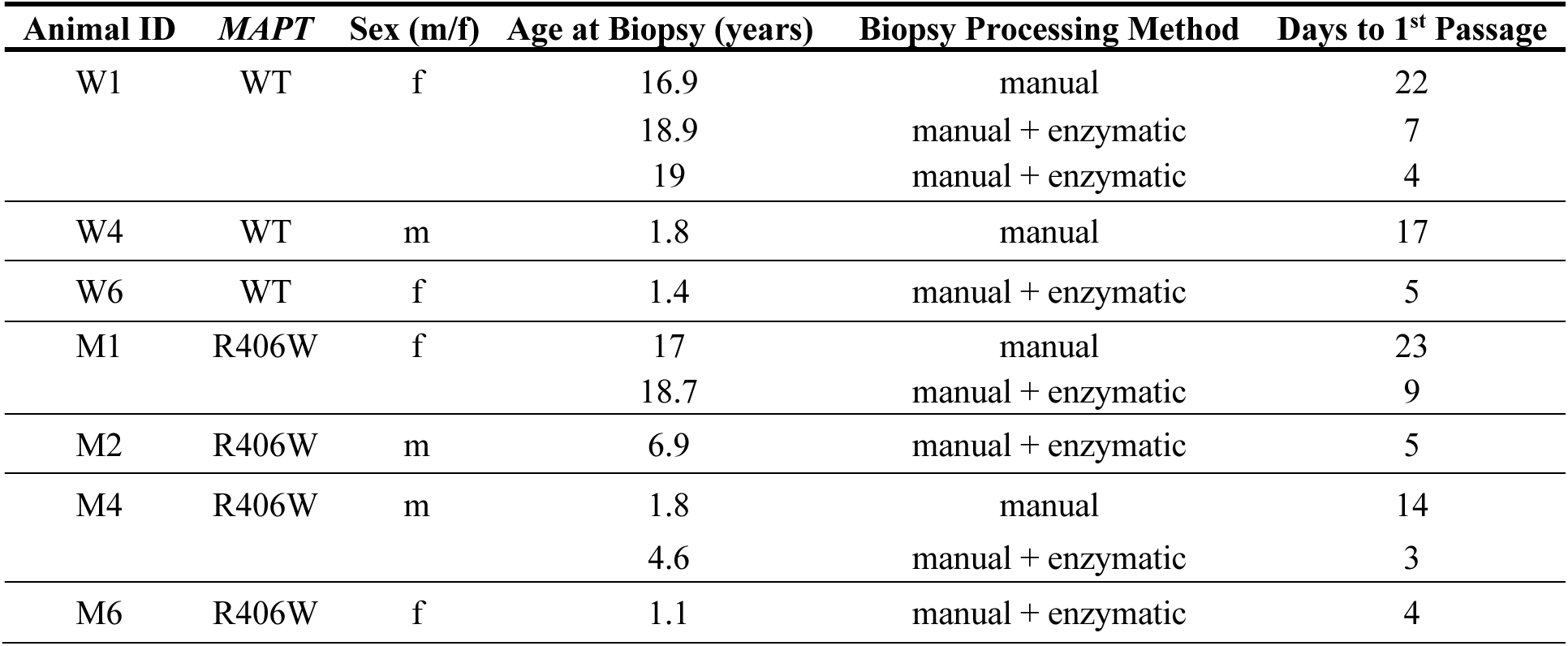
Animal demographics and assignment to skin biopsies. *Abbreviations: WT, wild type; m, male; f, female*.

Blood was obtained from *MAPT* R406W (n=2; 6.4-year-old male, 2.7-year-old female) and *MAPT* wild type (WT) rhesus macaques (n=2; 6.4-year-old male, 2.2-year-old female). Three animals underwent blood collection without sedation in a table-top restraint device, and one animal was under sedation (10 mg/kg ketamine and 0.2 mg/kg midazolam IM) for a separate procedure. A 2-3 mL blood sample was withdrawn from the femoral or saphenous vein into an EDTA tube using a vacutainer system.

### 2.3 Fibroblast Derivation from Skin Punch Biopsies

Two methods were tested to establish rhesus fibroblasts from skin punch biopsies: manual processing alone and manual plus enzymatic processing. For manual processing alone (n=4 biopsies), biopsy tissue was placed into a sterile 10 cm tissue culture dish, a scalpel was used to remove any remaining hair, and each biopsy was cut into 2-4 pieces. Biopsy pieces were then directly transferred to individual wells of a 6-well plate pre-coated with 0.1% gelatin (Sigma-Aldrich, cat. no. ES006B) and containing 1 mL rhesus fibroblast medium. For manual plus enzymatic processing (n=7 biopsies), the biopsy pieces resulting from manual processing were additionally enzymatically digested overnight in digestion medium with incubation at 37°C and 5% CO_2_. The digestions were vortexed briefly and the tissue pieces were removed and plated as described above. The remaining cell suspensions were centrifuged at 210 x g for 5 min to isolate the cell pellets. The cell pellets were resuspended in 1 mL fibroblast medium and then distributed to the 6-well plate containing the tissue pieces. Cells and tissue pieces were incubated at 37°C in 5% CO_2_. Media was exchanged after 24 h post-plating and thereafter exchanged every other day.

The fibroblasts were passaged when they were either (1) densely concentrated around the biopsy tissue and in need of redistribution or (2) >90% confluent when evenly distributed in a monolayer. Fibroblasts were first incubated with 0.25% trypsin (Gibco, cat. no. 15090046) for 5-10 min at 37°C. Cell pellets were isolated by centrifugation at 210 x g for 5 min, and cells were re-plated at a density of 1.5-2×10^5^ cells per well of a 6-well plate or approximately 1×10^6^ cells per T75 flask. Fibroblasts that had undergone up to four passages were either directly reprogrammed to RhiPSCs or cryopreserved at a density of 1-1.5×10^6^ cells per vial in CELLBANKER^®^ 1 (Amsbio, cat. no. 11910) until reprogramming.

### 2.4 Fibroblast Reprogramming to RhiPSCs

Non-integrating Sendai viral vectors and episomal (oriP/EBNA1) plasmids were evaluated for delivery of reprogramming factors to fibroblasts and generation of RhiPSCs.

#### 2.4.1 Sendai Virus Reprogramming

Three reprogramming trials with the CytoTune™-iPS 2.0 Sendai Reprogramming Kit (ThermoFisher Scientific, cat. no. A16517), were performed using one *MAPT* WT fibroblast line (W4) and one *MAPT* R406W (M4) fibroblast line in a 6-well plate format (**Table 2**). Variables assessed between reprogramming trials included the starting number of fibroblasts, viral titers, and different iPSC media formulations.

**Table 2:**
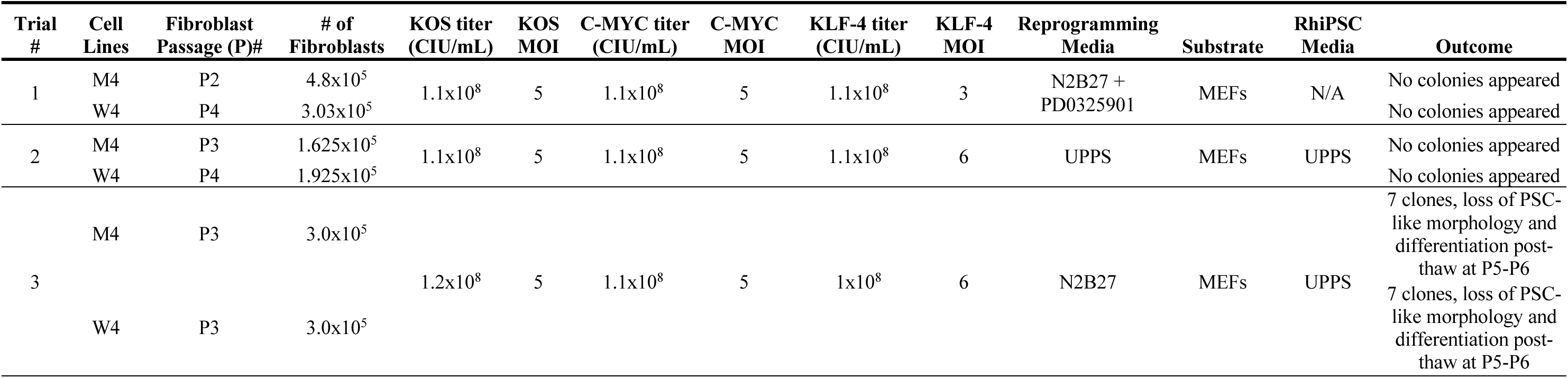
Reprogramming trials with Sendai viral vectors. *Abbreviations: CIU, cell infectious units; MEFs, mouse embryonic fibroblasts; N/A, not applicable; UPPS, Universal Primate Pluripotent stem cell medium*.

The first trial was performed following the recommended protocol for feeder-dependent fibroblast reprogramming by the CytoTune™-iPS 2.0 Sendai Reprogramming Kit. Briefly, on day 0, 2 wells/cell line of 3-5×10^5^ fibroblasts each were transduced with KOS, C-MYC, and KLF-4 reprogramming vectors with a multiplicity of infection (MOI) of 5:5:3, respectively. At 24 h post-transduction, the spent cell culture media were replaced with fresh fibroblast media. On day 2, cells were passaged at a 1:3 dilution to fresh gelatin-coated 6-well plates due to over-confluence. On day 7, the cells were passaged at a 1:6 dilution to mouse embryonic fibroblast (MEF)-coated (3.9×10^4^ MEFs/cm^2^) 6-well plates. The cells were switched to N2B27 medium supplemented with 0.5 µM PD0325901 on day 8 and maintained until day 21 post-transduction.

In the second trial, 2 wells/cell line of 1.5-2×10^5^ fibroblasts each were transduced with KOS, C-MYC, and KLF-4 reprogramming vectors with a MOI of 5:5:6, respectively. At 24 h post-transduction, the spent cell culture media were replaced with fresh fibroblast media. On day 2, the cells were passaged at a 1:6 dilution to MEF-coated (3.9×10^4^ MEFs/cm^2^) 6-well plates. On day 3, the cells were switched to Universal Primate Pluripotency Stem Cell (UPPS; (Stauske et al., 2020)) medium. The cells were maintained until day 28 post-transduction.

For the third trial, 2 wells/cell line of 3×10^5^ fibroblasts each were transduced with KOS, C-MYC, and KLF-4 reprogramming vectors at a MOI of 5:5:6, respectively. The reprogramming vectors were added directly to N2B27 medium without PD0325901 (Tao et al., 2021), rather than to the rhesus fibroblast medium as done in the other two trials. At 24 h post-transduction, cells were transferred to Growth Factor Reduced (GFR) Matrigel-coated (Corning Life Sciences, cat. no. 354230) 6-well plates and fed fresh N2B27 medium. On day 8, the cells were switched to UPPS medium. RhiPSC colonies that appeared were picked and transferred to MEF-coated plates (1.95×10^4^ MEFs/cm^2^) for clonal expansion and fed UPPS medium daily.

#### 2.4.2 Episomal Reprogramming

A total of twelve reprogramming trials were performed using electroporation of non-integrating oriP/EBNA1 plasmids to deliver reprogramming factors to fibroblasts (Yu et al., 2009). The plasmids were delivered to cells using a Neon Transfection system (ThermoFisher Scientific, model no. MPK5000) which has 24 fixed programs with unique electroporation settings (i.e., number of pulses (range 1-3), pulse voltage (range 850-1600V), and pulse width (range 10-40 ms) (**Supplementary Table 2**)). The modified variables included the number of fibroblasts transfected, electroporation conditions, amount of plasmid and combination of reprogramming factors, transfected cell plating density, and RhiPSC culture media (**Table 3**). In all reprogramming trials, RhiPSC colonies that appeared were manually picked to MEF-coated 24-well plates (3.9×10^4^ MEFs/cm^2^) to establish clonal lines and expand for further characterization and cryopreservation.

**Table 3:**
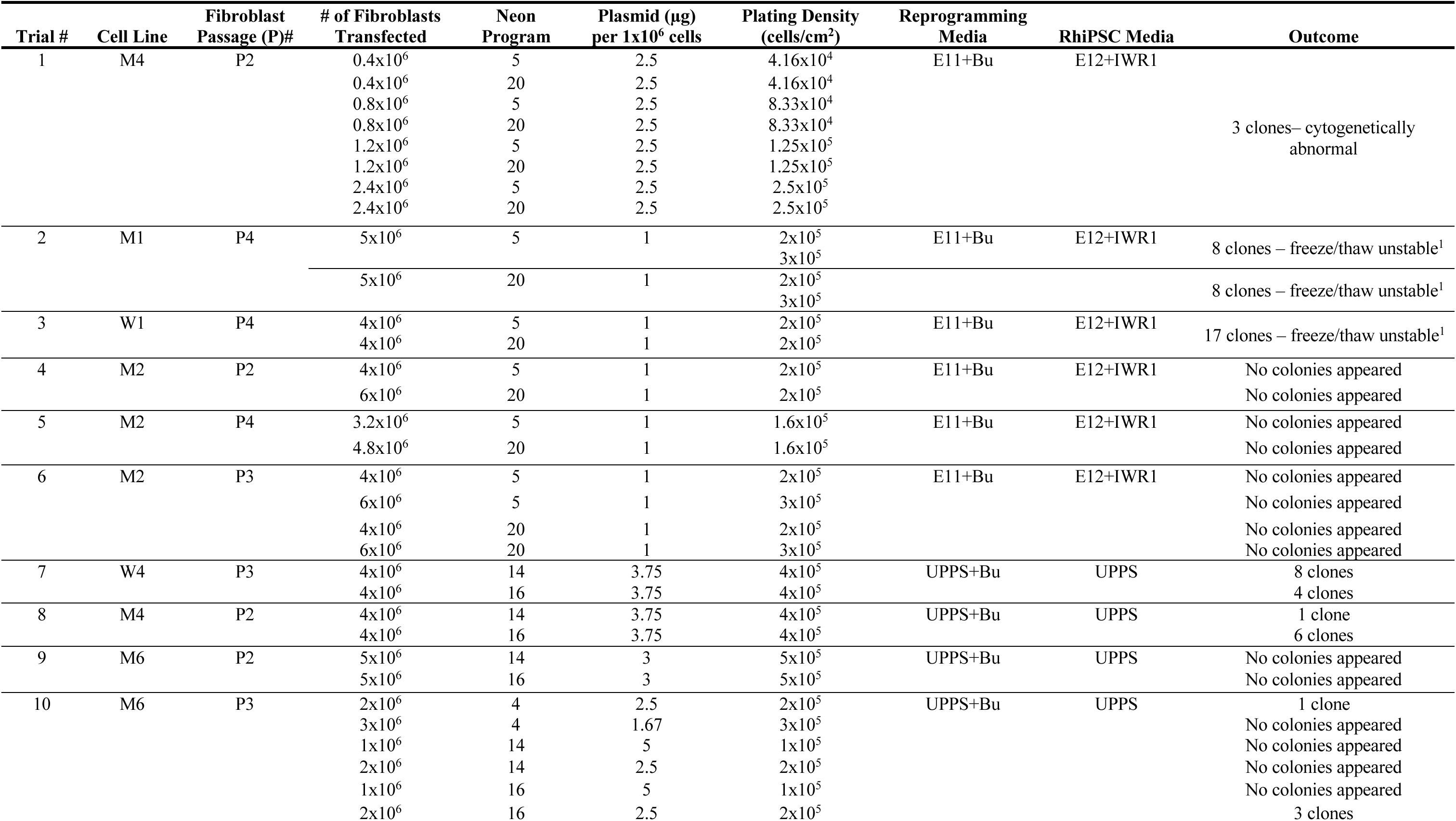
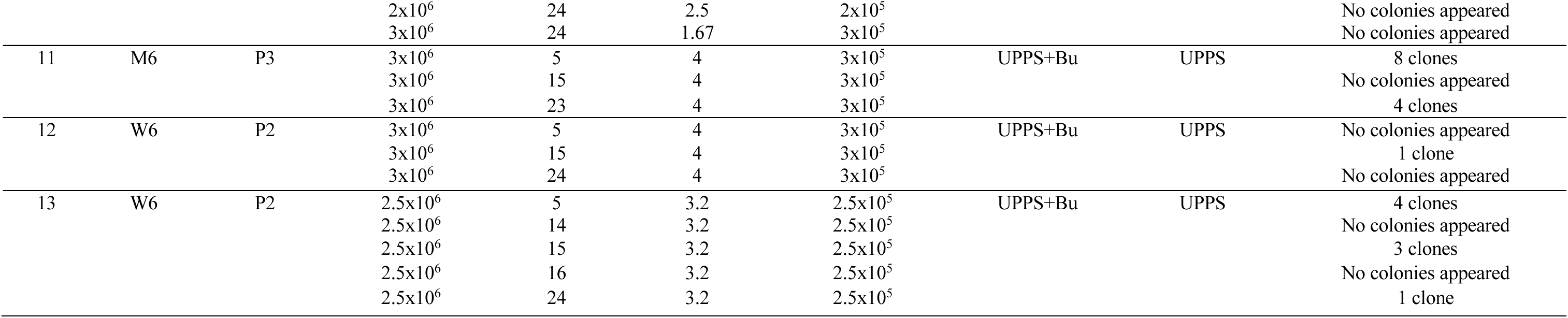
Reprogramming trials with oriP/EBNA1 episomal plasmids. Abbreviations: E11, Essential 11 medium; E12, Essential 12 medium; IWR1, inhibitor of Wnt response-1; MEFs, mouse embryonic fibroblasts; UPPS, Universal Primate Pluripotent stem cell medium; +Bu, plus sodium butyrate. ^1^RhiPSC Morphology not maintained post-thaw.

Trials 1-6 involved fibroblasts from animals M4, M1, W1, and M2 (**Table 3**). For these trials, fibroblast reprogramming was initiated on Neon programs #5 and #20 (chosen based on previous successful in-house cynomolgus macaque fibroblast reprogramming events; unpublished data) by electroporating cells with plasmids pEP4 E02S LM2L (Addgene, plasmid #20926), pEP4 E02S ET2K (Addgene, plasmid #20927), pEP4 E02S EN2K (Addgene, plasmid #20925) and pSimple-miRNA302/367 (Addgene, plasmid #98748), combined with EBNA1 mRNA *in vitro* transcribed from the template Sp6-EBNA1 (Addgene, plasmid #98749) (Chen et al., 2011; Howden et al., 2015; Howden et al., 2006; Yu et al., 2009) using the mMessage mMachine SP6 transcription kit (ThermoFisher Scientific, cat. no. AM1340). EBNA RNA was quantified by UV light absorbance (IMPLEN NanoPhotometer N50) and the length of the transcript (i.e., 1248 bp) was analyzed by denaturing gel electrophoresis. *In vitro* transcribed mRNA encoding a truncated version of the EBNA1 protein was included to enhance nuclear uptake of the oriP-containing reprogramming plasmids (Chen et al., 2011; Howden et al., 2006). In trial 1, transfected cells were plated at varying densities (range 4.16×10^4^-2.5×10^5^ cells/cm^2^) to Vitronectin-coated 6-well plates (50 μg/plate; **Table 3**). In subsequent trials, 2-3×10^6^ cells were plated onto Vitronectin-coated 10 cm tissue culture dishes (50 µg/dish) containing 10 mL of fibroblast medium immediately following electroporation. The cells were transitioned to Essential 11 (E11) medium supplemented with 100 μM sodium butyrate (E11+Bu) on day 3 post-electroporation and then to Essential 12 (E12) medium supplemented with the Wnt inhibitor IWR1 (E12+IWR1) on day 10 (Liang et al., 2010).

In trials 7-13, fibroblast reprogramming was initiated by electroporating fibroblasts from W4, M4, M6, and W6 (**Table 3**) with plasmids pEP4 E02S EN2L (Addgene, plasmid # 20922), pEP4 E02S ET2K, pEP4 E02S EM2K (Addgene, plasmid # 20923), pSimple-miRNA302/367, and EBNA1 mRNA. In trials 7-9, fibroblasts from W4, M4, or M6 were electroporated on Neon programs #14 and #16 (chosen based on previous successful in-house rhesus macaque fibroblast reprogramming events (Jacobo Lopez et al., 2022)). Subsequently, trials 10-13 for M6 and W6 fibroblasts were initiated on Neon programs selected following optimization of plasmid electroporation conditions (see Section 2.4.3); these included Neon programs #5, #14, #16, and #24 for both fibroblast lines, plus Neon programs #4 and #23 for line M6 and Neon program #15 for line W6. Cells were maintained in fibroblast medium until day 4 post-electroporation and then switched to UPPS medium containing 100 μM sodium butyrate (UPPS+Bu). Sodium butyrate was removed from the medium after seven days.

#### 2.4.3 Optimization of Plasmid Electroporation

To optimize plasmid electroporation, all 24 Neon Transfection(ThermoFisher Scientific; **Supplementary Table 2**) programs were evaluated in fibroblasts derived from M4, M6, W4 and W6 (two experimental replicates per fibroblast line). For each cell line, 1×10^5^ fibroblasts were resuspended in R buffer (10 μL/electroporation; ThermoFisher Scientific, cat. no. MPK1096) with enhanced Green Fluorescent Protein (eGFP) cloned into a pSimpleII plasmid (an OriP-containing plasmid; 0.5 μg/electroporation) as previously described (Hou et al., 2013; Howden et al., 2015) under the control of the elongation factor-1α promoter (EF1α) using a 10 μl Neon electroporation tip for each program. The cells were then seeded to individual wells of a 24-well plate (1 well per Neon program) pre-coated with Vitronectin (50 μg/plate). Cells were imaged at 24 and 48 h post-electroporation using a Nikon Eclipse Ti2 microscope with NIS Elements software version 6.10.02 (Nikon, Tokyo, Japan). Phase images were captured on a DS-Ri2 camera to assess fibroblast survival and confluency. eGFP fluorescence was imaged on a DS-Qi2 camera to assess plasmid uptake efficiency.

Cells were collected at 48 h post-electroporation to assess the percentage of live and eGFP positive cells via flow cytometric analysis using a MACSQuant Analyzer 10 Flow Cytometer (Miltenyi Biotec). Briefly, cells were lifted with Accutase (ThermoFisher Scientific, cat. no. A1110501), quenched with PBS (ThermoFisher Scientific, cat. no. BP3991) supplemented with 2% FBS (PBS+FBS; Peak Serum, cat. no. PSFB4) and spun at 300 x g for 5 min. Cells were stained with 0.25 μL Ghost Dye Violet 540 (Tonbo Bioscience, cat. no. 130879T100) viability dye for 20 min in PBS+FBS. The percentage of live cells expressing eGFP was calculated using FlowJo 10.10.0 software (BD Life Sciences). Briefly, the main cell population was selected from forward scatter (FSC) and side scatter (SSC). Single cells were then distinguished from doublets or aggregates using FSC-height and FSC-area from the main cell population gate. The viability dye Ghost Dye Violet 540 (Fisher Scientific, cat. no. 50-201-4118) was used to identify and exclude dead cells from the violet 405 nm laser 525/50 nm filter set and eGFP expressing positive cells were identified from the blue 488 nm laser 525/50 nm filter set.

### 2.5 RhiPSC Maintenance

RhiPSCs were maintained at 37°C and 5% CO_2_ in a humidified incubator and expanded on MEF-coated 6-well plates (3.9×10^4^ MEFs/cm^2^) in UPPS medium (2 mL/well) with daily media changes. Cells were passaged at a 1:3 dilution every 2–4 days by incubation with 0.5 mM EDTA in 1×PBS without calcium chloride or magnesium chloride for 5-10 min at 37°C. EDTA was then aspirated and cells were gently agitated using a pipette containing UPPS medium (1 mL/well).

For cryopreservation, cells were grown to ∼80% confluency, then incubated with 0.5 mM EDTA as described above. EDTA was aspirated, then cells were gently agitated using a pipette containing Cryostor CS10 (1 mL/well of a 6-well plate; STEMCELL Technologies, cat. no. 72562). Cell suspensions from individual wells were transferred to single cryovials, respectively, and immediately placed in a controlled rate freezing container (ThermoFisher Scientific, cat. no. 5100-0001) in a -80°C freezer. After 24 h, the cryovials were transferred to liquid nitrogen for long-term storage. To thaw RhiPSCs, cryovials were warmed at 37°C. The cell suspension was then diluted in UPPS medium and centrifuged at 500 x g for 5 min. Cells were resuspended in UPPS medium and plated to 6-well MEF-coated plates (1 cryovial/well).

#### 2.5.1 Optimization of RhiPSC Culture Conditions

To identify optimal passage conditions, a clonal RhiPSC line from each of the four animals was plated to different matrices with or without the supplementation of supporting reagents. MEF-coated (3.9×10^4^ MEFs/cm^2^), GFR Matrigel-coated (0.5 mg/plate; Corning Life Sciences, cat. no. 354230), or Vitronectin-coated (50 μg/plate) 6-well plates were prepared. Prior to the addition of cells, 2 mL of either UPPS medium, UPPS medium plus 10 uM Y-27632 (RHO/ROCK pathway inhibitor; Tocris, cat. no. 1254), or UPPS medium plus 1XCultureCEPT Supplement (chroman 1, emricasan, polyamines, trans-ISRIB; CEPT; ThermoFisher Scientific, cat. no. A56799; (Chen et al., 2021)) were added to individual wells (one 6-well plate per condition of matrix + media per cell line). RhiPSCs at P14-P23 were dissociated with Accutase for 5 min or until fully detached from the plate, neutralized with equal volumes of Complete Medium, counted, washed, and diluted to 50,000 cells/mL. Suspensions of 100 μL (5,000 cells) were added into each well. Cells were incubated at 37°C and 5% CO_2_. After 24 h, UPPS medium containing Y-27632 or CEPT was replaced with UPPS medium without supplementation. Media was exchanged daily. After 5 days, colonies were stained with a Vector Blue Alkaline Phosphatase Substrate kit (Vector Laboratories, cat. no. SK-5300) as described in Section 2.6.1. The wells were then imaged and the number of colonies per well were counted. Percent cloning efficiency was calculated by dividing the colony counts for each well by the initial 5,000 seeded cells and then multiplied by 100. The number of colonies was averaged over 6 wells for each condition (matrix + media) per cell line. Data were analyzed by two-way repeated-measures ANOVA, with matrix and media as within-subject factors in GraphPad Prism v10.6.1. Post hoc comparisons were performed using Tukey’s multiple comparisons test, and significance threshold of p < 0.05 was applied.

### 2.6 RhiPSC Characterization

Following ISSCR standards (Ludwig et al., 2023), RhiPSCs were characterized by morphology, alkaline phosphatase staining, gene expression, immunocytochemistry, flow cytometry, karyotype, genotype, cell authentication, mycoplasma and viral screening and teratoma formation. See primer sequences in **Supplementary Table 3** and antibody information in **Supplementary Table 4**. All RhiPSC characterization experiments were performed on cells that had undergone at least one freeze/thaw cycle.

#### 2.6.1 Alkaline Phosphatase Staining, Immunocytochemistry, and Microscopy

Alkaline phosphatase (AP) staining was performed on RhiPSCs between P10-P22 using the Vector® Blue Substrate Kit (Vector Laboratories, cat. no. SK-500). Briefly, cells were passaged onto MEF-coated 6-well plates as described in Section 2.5 and AP staining was performed once the cells reached 50-80% confluency. The AP working solution was prepared according to the manufacturer’s directions. Prior to staining, media was removed, and cells were washed with 2 mL 1XPBS. Cells were stained by adding 2 mL of the AP working solution and incubated at 37°C for 20-30 mins. AP working solution was then removed, cells were washed with PBS to remove AP working solution, and imaged (D’Souza et al., 2016).

RhiPSCs between P11-P23 were seeded and cultured on Vitronectin-coated (50 μg/plate) glass-bottom culture dishes (MatTek, cat. no. P06G-0-20-F). Cells were fixed with 4% PFA for 15 min at 37°C followed by permeabilization with cold 90% methanol (10% PBS+FBS, -20°C) for 30 min. Samples were washed with PBS+FBS followed by staining with primary antibodies OCT3/4, SOX2, and NANOG diluted in PBS+FBS at 4°C overnight. The next day, cells were washed with PBS+FBS prior to incubation with AlexaFluor secondary antibodies diluted in PBS+FBS for 30 min and then counterstained with 4’,6-diamidino-2-phenylindole (DAPI) for 15 min at room temperature, covered from light. Cells were then washed with PBS+FBS and then imaged on a Nikon A1 confocal microscope with NIS Elements software version 5.41.02 (Nikon, Tokyo, Japan). Negative controls were processed in parallel by omitting primary antibodies during the procedure.

#### 2.6.2 RNA Isolation, Reverse Transcription (RT), and RT-PCR Analysis

RhiPSCs between P12-P19 were directly lysed with RLT Plus lysis buffer (Qiagen, cat. no. 1053393) and stored at -20°C or -80°C for later RNA processing. Total RNA was purified from the RLT Plus lysates using the RNeasy Plus Micro Kit (Qiagen, cat. no. 74034) according to the manufacturer’s instructions using genomic DNA (gDNA) removal columns. Purified RNA was reverse transcribed with the SuperScript III First-Strand cDNA Synthesis kit (ThermoFisher Scientific, cat. no. 18080051).

RT-PCR with primers specific to *POU5F1*, *NANOG*, *SOX2*, *KLF4*, *LIN28A*, and *MYC* were used to confirm the pluripotency of the RhiPSCs. A Platinum II Hot-Start DNA polymerase kit (ThermoFisher Scientific, cat. no. 14966001) was combined with gene specific primers (e.g., *POU5F1, NANOG, LIN28A, MYCT, ACTB*) and 100-200 ng of cDNA. A 20 µl reaction was performed using the 2-step cycling conditions as outlined in the manufacturer’s protocol. To amplify *SOX2* and *KLF4*, 100-200 ng of cDNA was combined with GoTaq G2 Hot Start master mix (Promega, cat no. M7422) and cycled using touchdown cycling conditions. RNA from a previously described rhesus embryonic stem cell line (RhESC: r420; (Thomson et al., 1995)) was included as a positive control cell line. *Beta-actin* (*ACTB*) was amplified as a positive control reference gene for each RhiPSC line, and a no-template negative control reaction for each primer were run in parallel. The PCR amplicons were run on 2.0% agarose gel at 110V for 45-55 min to verify amplification. A 100 bp DNA ladder (Promega, cat. no. g2101) was used to confirm the product size.

#### 2.6.3 Flow Cytometry

RhiPSCs between P16-P25 were singularized with Accutase, quenched with PBS+FBS, and spun at 300 x g for 5 min. Cell pellets were fixed with 4% paraformaldehyde (PFA) for 10 min at 37°C followed by permeabilization with cold 90% methanol and 10% PBS+FBS at -20°C for 30 min. Samples were washed with PBS+FBS followed by staining with unconjugated primary antibodies OCT3/4, SOX2, and NANOG (**Supplementary Table 4**) diluted in PBS+FBS at 4°C overnight. Cells were then washed with PBS+FBS buffer before incubation with secondary antibodies diluted in PBS+FBS for 30 min. The stained cells were washed with PBS+FBS and cytometric analysis was performed on MACSQuant X Flow Cytometer (Miltenyi Biotec). A single cell population was identified as described in Section 2.4.3, then allophycocyanin (APC) positive cells were detected with the red 640 nm laser 655-730 nm filter set.

#### 2.6.4 DNA Isolation and PCR

Total genomic DNA (gDNA) was extracted using the DNeasy Blood and Tissue Kit (Qiagen, cat. no. 69504) from ∼1×10^6^ cells.

To confirm loss of reprogramming episomal plasmids, PCR for *EBNA1* and oriP was performed using gDNA extracted from RhiPSCs at P14-P18. PCR reactions were conducted with GoTaq DNA Polymerase (Promega, cat. no. M7122) using 50 ng of gDNA as a template. gDNA from rhESCs (r420) served as a negative control while the EM2K reprogramming plasmid was used as a positive control. The reactions were cycled as follows: 94°C for 5 min, (94°C 15 s, 60°C 30 s, 68°C 1 min) x 25 cycles, 68°C for 7 min. PCR products were loaded to 2% agarose gels and ran at 70V for 90-120 min. The TrackIt 100bp DNA Ladder (ThermoFisher Scientific, cat. no. 10488058) was used to confirm product size.

To confirm maintenance of the *MAPT* genotype, gDNA collected from RhiPSCs at P13-P19 was assessed using standard PCR with primers specific to the *MAPT* R406W region. A 50 µl reaction containing Platinum II Hot-Start PCR Master mix (ThermoFisher Scientific, cat. no. 1400013) and 100 ng gDNA was assembled and cycled using the 3-step cycling protocol recommended by the manufacturer. The PCR amplicons were run on a 1.5% agarose gel at 80V for 180 min to confirm amplification. The QIAquick PCR Purification Kit (Qiagen, cat. no. 28104) was used for purification of the PCR products, following the manufacturer’s instructions. The purified PCR products were then diluted to 30 ng/mL and submitted for amplicon sequencing at the University of Wisconsin-Madison Biotechnology Center. Sequencing reads were aligned to the Mmul10 reference genome.

#### 2.6.5 Cell Line Authentication

Short tandem repeat (STR) analysis was performed and interpreted by the Veterinary Genetics Laboratory at the University of California-Davis to authenticate all eight RhiPSC lines (P16-P22). Blood samples served as sources of reference DNA.

#### 2.6.6 Mycoplasma and Viral Screening

Mycoplasma PCR screening was performed by WiCell (Madison, WI). RhiPSCs at P11-P20 were cultured for 24 h as described in Section 2.5 and 2 mL of spent media was submitted for evaluation. Routine viral screening was performed by Charles River Research Animal Diagnostic Services (Wilmington, MA). Briefly, 1-4×10^6^ fibroblasts were spun down at 300 x g for 5 min. The supernatant was aspirated and cell pellets were frozen at -80°C. Cell pellets were submitted for PCR detection of Herpes B virus (HBV), Simian Immunodeficiency virus (SIV), Simian T-Lymphotropic virus (STLV), and Simian Retrovirus (SRV).

#### 2.6.7 Karyotyping

Standard g-banded karyotyping was performed on RhiPSCs at P10-P18 and interpreted by Cell Line Genetics (Madison, WI). Briefly, cells were passaged into a MEF-coated (3.9×10^4^ MEFs/cm^2^) T-25 culture flask. Cells were submitted to Cell Line Genetics when they reached 60-70% confluency with large colonies that could be seen macroscopically.

#### 2.6.8 Teratoma Assays

RhiPSCs (∼8-10×10^6^ cells) at P16-P22 were suspended in a 350 μL solution of 50% UPPS medium and 50% GFR Matrigel. 100 μL suspensions were injected into the left hind leg muscle of female NCG-X mice (Charles River; n=3 mice/RhiPSC line) using a 27G needle. Mice were monitored daily by animal caretaking staff and palpated weekly for teratoma formation. Once tumors exceeded 2 cm in diameter, mice were euthanized by CO_2_ inhalation and teratomas collected postmortem. Teratomas were typically harvested eight to twelve weeks post-injection and fixed in 4% PFA for 48-72 h. The teratomas were placed in cassettes, dehydrated in 70% ethanol for 24 h, and sent to the School of Veterinary Medicine at UW-Madison for paraffin embedding and hematoxylin & eosin (H&E) staining. H&E-stained tissue sections were evaluated by board-certified pathologists (HAS, PB) and imaged on a Nikon Eclipse Ti2 microscope with a DS-Ri2 camera using NIS Elements software version 6.10.02 (Nikon, Tokyo, Japan).

### 2.7 RhNPC Derivation

RhiPSCs (1 line per animal) were differentiated to RhNPCs over 21 days using a monolayer method (Karch et al., 2019). On day 0 of differentiation, RhiPSCs were singularized by incubation with Accutase for 5-10 min at 37°C. Cell dissociations were quenched using Complete Medium, then centrifuged at 500 x g for 5 min. Cell pellets were resuspended in Neural Induction Medium (NIM) supplemented with 10 μM Y-27632. RhiPSCs were plated to 6-well plates pre-coated with Poly-L-Ornithine (PLO; Sigma-Aldrich, cat. no. P4957) and mouse laminin (Sigma-Aldrich, cat. no. L2020; 10 mg/mL) at a density of 2×10^6^ cells per well. Cells were incubated at 37°C and 6% CO_2_; NIM medium (lacking Y-27632) was exchanged daily.

RhNPCs were passaged every 7 days. At each passage, cells were singularized in Accutase as described above, then the cell suspension was centrifuged at 170 x g for 5 min. Cells were resuspended in NIM supplemented with 10 μM Y-27632 and then plated to PLO and mouse laminin pre-coated 6-well plates at a density of 2×10^6^ cells per well. RhNPCs were collected on day 21 of differentiation for characterization of NPC marker expression or cryopreserved at a density of 1-2×10^6^ cells per vial in 500 μL NIM and 500 μL Neural Freezing Medium.

### 2.8 RhNPC Immunocytochemistry and Microscopy

For immunocytochemical staining of neuronal markers, day 21-differentiated RhNPCs were lifted and replated to PLO/laminin-coated, glass-bottom 6-well plates (MatTek, cat. no. P06G-0-20-F) at a density of 0.5×10^6^ cells per well in NIM supplemented with 10 μM Y-27632. After 24 h of incubation at 37°C, cells were fixed, stained, and imaged as described in Section 2.6.1. Primary antibodies included PAX6 and NESTIN (**Supplementary Table 4**). Negative controls were processed in parallel by omitting primary antibodies during the procedures. Z-stack immunofluorescence images were captured on a Nikon A1 confocal microscope and maximum intensity projection images were rendered using NIS Elements software version 5.41.02 (Nikon, Tokyo, Japan). Look up tables (LUTs) were adjusted to enhance signal.

## 3.0 Results

### 3.1 Fibroblast Derivation from Skin Punch Biopsies

Manual tissue processing (i.e., mincing of dermal biopsy tissue), typically used for processing of human skin biopsies, was first evaluated for rhesus fibroblast derivation. Manual tissue processing alone resulted in limited initial rhesus fibroblast outgrowth that led to a longer propagation time. The average time elapsed between plating tissue biopsy pieces, and the first passage (P0 to P1) was 19 days (range 14-23 days; n=4 biopsies; **Table 1**). The cells were passaged until ∼P4 to obtain enough cells for cryopreservation or reprogramming. The fibroblasts derived from *MAPT* WT rhesus presented typical, elongated morphology regardless of passage number (**Supplementary Figure 1A**). In comparison, fibroblasts derived from *MAPT* R406W rhesus starting at P3-P4 became enlarged, multinucleated, and detached from the culture vessel surface (**Supplementary Figure 1B**).

Overnight enzymatic digestion with Collagenase type I and DNase I resulted in more fibroblast growth and yield with a shorter propagation time. The average period elapsed between plating tissue biopsy pieces, and the first passage (P0 to P1) was 5.3 days (range 3-9 days; n=7 biopsies; **Table 1**). The addition of the overnight enzymatic digestion allowed for collection of low passage (≤P2) fibroblasts for cryopreservation or reprogramming. At ≤P2, fibroblasts derived from *MAPT* WT and R406W rhesus were morphologically identical and viable. Low passage (≤P2) fibroblast lines were ultimately established using the manual plus enzymatic digestion method from 3 *MAPT* WT and 4 *MAPT* R406W rhesus **(Table 1)**. It should be noted that regardless of processing method and animal age at biopsy collection, *MAPT* R406W-derived fibroblasts at ≥P3 were morphologically abnormal and not viable.

### 3.2 Fibroblast Reprogramming to RhiPSCs

#### 3.2.1 Sendai Virus Reprogramming

Sendai viral vectors were first tested for reprogramming rhesus fibroblasts as it is the most widely utilized method for both hiPSC and RhiPSC derivation (Jocher et al., 2024; Lara et al., 2023; Pozner et al., 2025; Tao et al., 2021; Zhao et al., 2018). Three Sendai virus reprogramming trials were performed with the CytoTune™-iPS 2.0 Sendai Reprogramming Kit; all had poor outcomes. The modified variables between the Sendai reprogramming trials included the number of fibroblasts transduced, viral multiplicity of infection (MOI), and cell culture media; these variables and trial outcomes are reported in **Table 2**.

The first trial followed the suggested protocol for feeder-dependent reprogramming of fibroblasts provided by the kit, including the recommended MOI and the suggested number of fibroblasts (i.e., ∼3.0×10^5^ cells/well; W4). More fibroblasts were transduced for line M4 (i.e., 4.8×10^5^ cells/well) to ensure a sufficient number of proliferating fibroblasts given the previous observation of cell death in the *MAPT* R406W fibroblasts (**Supplementary Figure 1B**). Within the first week following viral transduction, excessive fibroblast proliferation was observed; this necessitated a passage at a 1:3 dilution on day 2 post-transduction and a passage at a 1:6 dilution when cells were to be transferred to MEFs on day 7. The transduced cells were switched from rhesus fibroblast medium to N2B27 medium supplemented with PD0325901 on day 8, yet excessive fibroblast proliferation still occurred. On day 10, cells were passaged to new MEF-coated 6-well plates at dilutions of 1:12 or 1:20 due to over-confluence. By day 21, cells were overly confluent and still presented mostly fibroblast-like morphology; no RhiPSC colonies were observed. Cells lifted from the plate and were discarded.

For the second trial, the number of starting fibroblasts for transduction was decreased to 1.925×10^5^ cells/well for W4 and 1.625×10^5^ cells/well for M4. In addition, the MOI of the CytoTune 2.0 hKlf4 viral vector was increased, the RhiPSC medium was switched to UPPS medium as it was previously shown to support macaque iPSCs (Stauske et al., 2020), and the transduced cells were transferred to the MEF feeders 5 days earlier than in the first reprogramming trial. Despite starting with fewer fibroblasts, excessive proliferation was observed by 24-h post-transduction, again necessitating a passage at a 1:6 dilution in which cells were plated to MEFs on day 2. Between days 3 and 28, excessive fibroblast proliferation was observed, requiring passages at 1:20 or 1:40 dilutions every ∼5 days. Some wells became overly confluent and lifted from the plate, and no RhiPSC colonies were observed by day 28 post-transduction in the remaining wells. Overall, the cells again retained mostly fibroblast-like morphology, with excessive proliferation that exceeded what was observed during the first reprogramming trial.

The third trial was performed with the suggested number of fibroblasts recommended by the CytoTune™ kit (i.e., 3.0×10^5^ cells/well) and the viral concentrations used in trial 2, however, the Sendai viral vectors were introduced to fibroblasts in N2B27 medium instead of rhesus fibroblast medium, as it has previously supported RhiPSC derivation (Tao et al., 2021). The transduced cells were maintained in the N2B27 medium for one week, then were switched to UPPS medium on day 8 post-transduction. Cell colonies appeared on day 25 and 29 for the W4- and M4-derived cell lines, respectively. Seven colonies were manually isolated from each line for clonal expansion, however, for both RhiPSC lines, stem cell morphology could not be maintained after 5-6 passages and excessive differentiation was observed.

#### 3.2.2 Episomal Reprogramming and Electroporation Optimization

A total of thirteen trials were undertaken with oriP/EBNA1-based episomal plasmids to reprogram seven rhesus fibroblast lines into transgene-free RhiPSCs (**Table 3**), an approach previously reported to successfully derive human and cynomolgus macaque iPSCs (Baik et al., 2023; D’Souza et al., 2016; Yu et al., 2009). The modified variables included the number of fibroblasts transfected, electroporation conditions (i.e., Neon program), plasmid combination and amount, transfected cell plating density, and cell media.

Trials 1-6 utilized Neon programs #5 and #20 based on previous in-house reprogramming of cynomolgus macaque fibroblasts (unpublished data). RhiPSCs were derived in E11+Bu medium and expanded in E12+IWR1 medium. Trial 1 demonstrated that plating densities of 1.25-2.5×10^5^ transfected fibroblasts per cm^2^ of culture vessel best supported RhiPSC colony appearance. Thus, plating densities were ≥1.0×10^5^ transfected fibroblasts per cm^2^ of culture vessel in all subsequent trials (with slight variations dependent on the chosen electroporation program and individual cell lines). Trials 1-3 resulted in the establishment of RhiPSC lines from M1, W1, and M4 (**Table 3**). Trials 4-6 attempted to reprogram fibroblasts from M2 without success. The three RhiPSC lines generated from trials 1-3 expressed pluripotency proteins OCT4, SOX2, and NANOG but displayed aberrant morphology; colonies were neither compact nor formed distinct borders (**Supplementary Figure 2A**). Furthermore, the RhiPSC line generated from animal M4 (trial 1) was cytogenetically abnormal (**Supplementary Figure 2B**). The RhiPSCs generated from animals M1 and W1 (trials 2 and 3, respectively) could not be reestablished after a freeze/thaw cycle and therefore were not karyotyped.

For reprogramming trials 7-9, E12 medium was replaced with UPPS medium for RhiPSC derivation, clonal expansion, and cell maintenance. Neon electroporation programs #14 and #16, selected based on previous successful in-house rhesus macaque fibroblast reprogramming to iPSCs (Jacobo Lopez et al., 2022), were tested and the reprogramming plasmid combination was modified (i.e., LM2L was replaced by EN2L and EN2K was replaced by EM2K). These changes led to successful derivation of RhiPSC lines from W4 (12 clones) and M4 (7 clones). Two clonal lines from each subject (W4, M4) were expanded and characterized according to ISSCR standards (see section 3.4). With the switch to UPPS medium, all four cell lines maintained pluripotency and normal karyotype over prolonged culturing. RhiPSC colonies from M6 were not generated (trial 9, **Table 3**).

Due to unsuccessful attempts at generating RhiPSCs from M6 in trials 4 and 9, we hypothesized that individual rhesus fibroblast lines may require optimization of the electroporation program for successful reprogramming. Considering reprogramming efficiency as a balance between plasmid uptake and cell survival after electroporation, the 24 Neon programs were evaluated across four fibroblast lines (M4, W4, M6, W6). The experiment was repeated twice for each fibroblast line, and the percent live cells and percent eGFP-positive cells at 48 h post-electroporation were quantified (**Figure 2A,C; Supplementary Figure 3A,C; Supplementary Table 2**). Data were visually inspected to select the programs that resulted in relatively high eGFP expression and cell survival. Overall, all four fibroblast lines had slight variations in the electroporation programs that resulted in the highest eGFP expression and post-electroporation cell survival, yet many of the same programs appeared as optimal across fibroblast lines (e.g., program #16 consistently resulted in high eGFP expression; **Figure 2B,D; Supplementary Figure 3B,D; Supplementary Table 2**). Programs considered best for fibroblast reprogramming included #5, #14, #15, #16, #20, #23, and #24.

**Figure 2.**
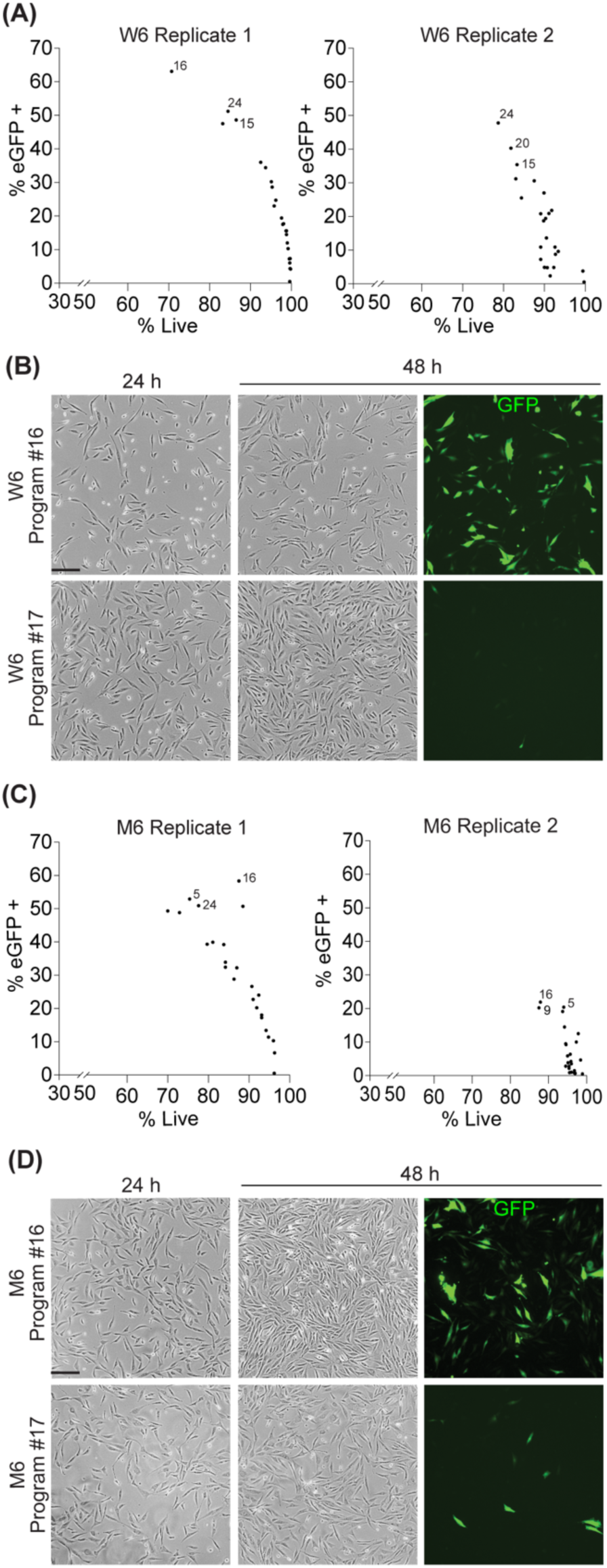
Optimization of electroporation efficiency for fibroblast lines W6 and M6 using the Neon Transfection system. The percentage of eGFP-expressing cells (% eGFP+; y-axis) were plotted against the percentage of live cells (% live; x-axis) for W6 (A) and M6 (C); individual dots represent data from each of the 24 different Neon programs. (B, D) Representative phase contrast images of fibroblasts lines W6 (B) and M6 (D) demonstrating post-electroporation cell survival at 24 and 48 h on Neon program #16 (an optimal program) and program #17 (a suboptimal program). Corresponding immunofluorescence images were taken 48 h post-electroporation to demonstrate the eGFP+ population resulting from Neon programs #16 and #17. Scale bars, 100 µm.

Reprogramming trials 10-13 were performed using the optimal programs for each respective fibroblast line. In trials 10 and 11, RhiPSCs were successfully derived from M6 fibroblasts using programs #5 (8 clones), #16 (3 clones), and #23 (4 clones). In trials 12 and 13, W6 RhiPSCs were generated with programs #5 (4 clones), #15 (4 clones), and #24 (1 clone). Two clonal lines from each subject (W6, M6) were expanded and characterized according to ISSCR standards, and all four cell lines maintained pluripotency and normal karyotype over prolonged culturing.

### 3.3 Optimization of RhiPSC Culture Conditions

Although the use of MEFs is usually reported in nonhuman primate (NHP) cell studies, feeder-free options such as Matrigel and Vitronectin are commonly used for hiPSCs. Thus, we compared the effect of the three matrices on RhiPSC clonal expansion. In addition, CEPT or Y-27632 were added to the UPPS medium to determine which supplement best supports RhiPSC self-renewal. Five days after RhiPSCs were seeded onto MEFs, typical compacted colonies with defined borders were observed. In contrast, RhiPSC colonies formed on Matrigel- and Vitronectin-coated plates appeared flatter with less distinct boundaries. Colonies that formed on Vitronectin-coated plates also tended to be smaller (**Figure 3A**). Experiments in 4 cell lines (n=2 *MAPT* WT; n=4 *MAPT* R406W) testing the three matrices with either CEPT, Y-27632 or no supplement, confirmed that both the growth matrix (p = 0.065, 43.8% variation) and supplement (p = 0.014, 27.7% variation) significantly impacted cloning efficiency. Individual subjects showed different trends in general cloning efficiency with 16% of the variability found to be subject-related. Overall, the combination of MEFs with CEPT supplementation was identified as the most optimal for clonal expansion (**Figure 3B**).

**Figure 3.**
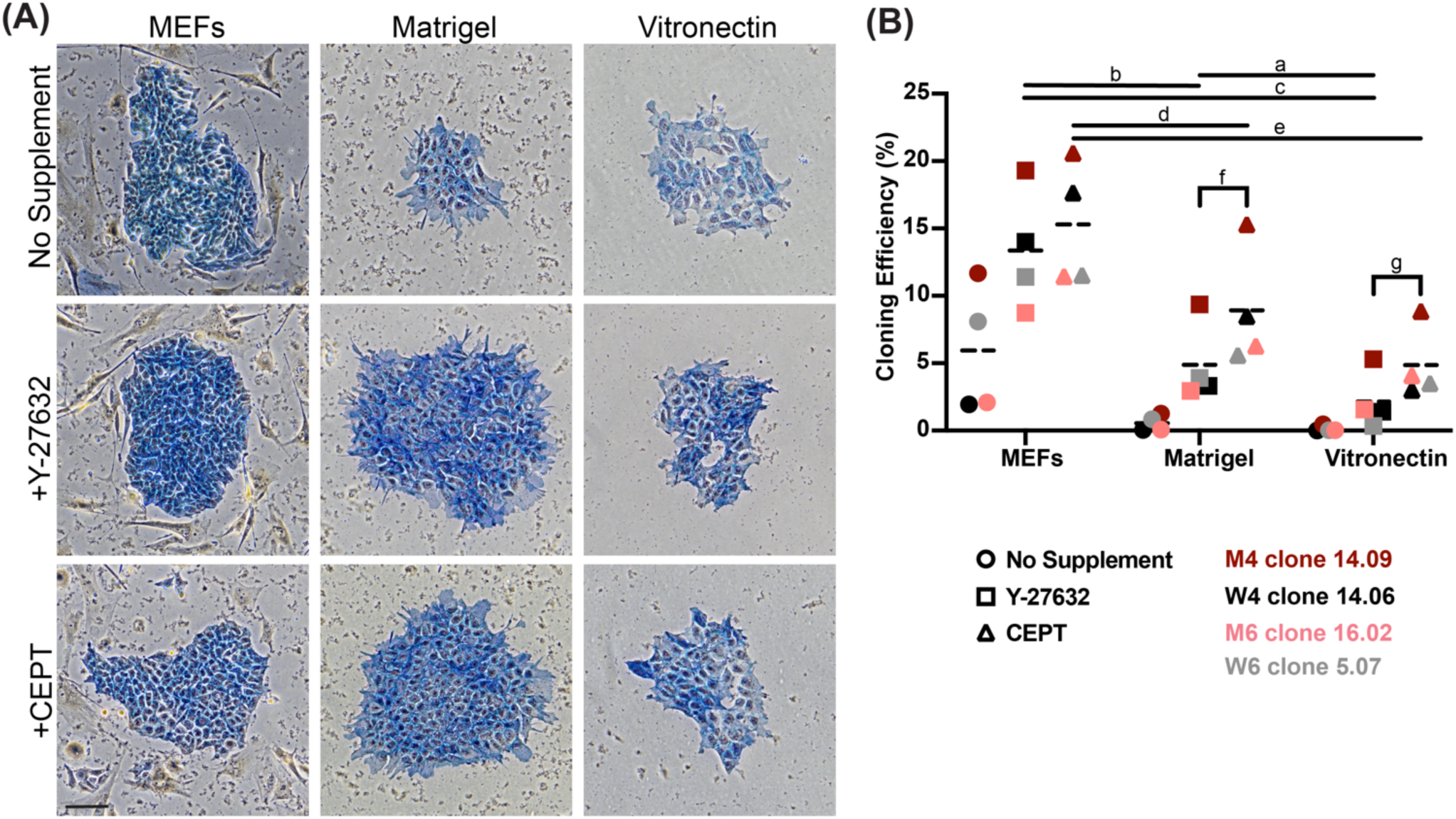
Effect of RhiPSC culture conditions on cloning efficiency. (A) Representative images of alkaline phosphatase-stained colonies from RhiPSC line M6 clone 16.02 on mouse embryonic fibroblasts (MEFs), Matrigel, or Vitronectin in UPPS medium with either no supplement, Y-27632, or CEPT. Scale bar, 100 µm. (B) Graph depicting percent (%) cloning efficiency (y-axis) of alkaline phosphatase-positive colonies cultured on different matrices (x-axis) in UPPS medium with no supplement (circles), Y-27632 (squares), or CEPT (triangles). Individual data points correspond to the mean (dashed lines) % cloning efficiency of RhiPSC colonies counted among six technical replicates per cell line (M4 clone 14.09, red; M6 clone 16.02, pink; W4 clone 14.06, black; W6 clone 5.07, grey). Black brackets indicate statistically significant differences between two supplement types within a substrate group; black lines indicate statistically significant differences between two substrates with the same supplement. p < 0.05 is considered statistically significant. Tukey’s multiple comparisons post-hoc p-values: a=0.0488, b=0.0098, c=0.0097, d=0.0133, e=0.0177, f=0.0483, g=0.0166.

### 3.4 RhiPSC Characterization and Maintenance

Eight RhiPSC lines were derived from 4 individual rhesus macaques: W6 (**Figure 4 & Supplementary Figure 4**), M6 (**Figure 5 & Supplementary Figure 5**), W4 (**Supplementary Figures 6, 7),** and M4 (**Supplementary Figures 8, 9**). Clonal lines were established by picking individual colonies and expanding them further for characterization; two clones per subject were fully characterized according to ISSCR standards (Ludwig et al., 2023).

**Figure 4.**
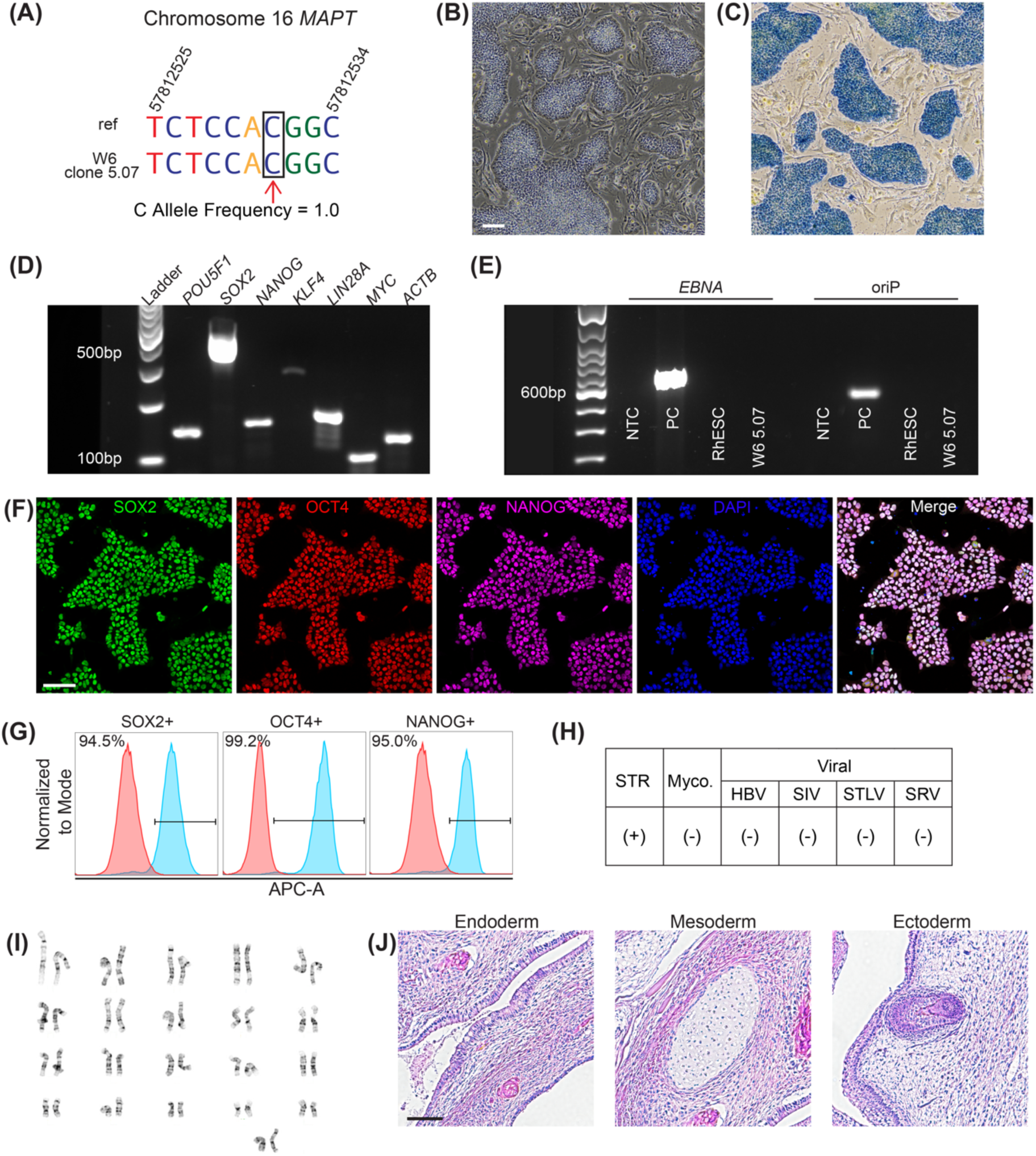
Characterization of RhiPSC line W6 clone 5.07. (A) Amplicon sequencing of *MAPT* R406W with wild-type C allele indicated in the black box with a red arrow compared to reference genome (ref). (B) Phase contrast image of colonies grown on MEFs and (C) positive alkaline phosphatase staining; scale bar, 100 µm. (D) RT-PCR detection of endogenous pluripotent mRNAs *POU5F1*, *SOX2*, *NANOG*, *KLF4*, *LIN28*, and *MYC*. *ACTB* was used as a reference gene. (E) Amplification of EBNA and oriP in RhiPSCs at P16, no template control (NTC), and the rhesus embryonic stem cell line r420 (RhESC). Plasmid EM2K was used as a positive control (PC) for the detection of EBNA and oriP. (F) Positive immunofluorescence staining for pluripotency proteins SOX2 (green), OCT4 (red), and NANOG (far red), colocalized with DAPI (blue). Scale bar, 100 µm. (G) Flow cytometry-detected APC-A+ cells were 94.5% positive for SOX2, 99.2% positive for OCT4, and 95% positive for NANOG. (H) Short tandem repeat (STR) analysis confirmed (+) RhiPSC line identity. Mycoplasma (Myco.), Herpes B Virus (HBV), Simian Immunodeficiency Virus (SIV), Simian T-Lymphotropic Virus (STLV), and Simian Retrovirus (SRV) were not detected (-). (I) Normal 42,XX karyotype at P12. (J) H&E staining of a teratoma containing endoderm, mesoderm, and ectoderm. Scale bar, 100 µm.

**Figure 5.**
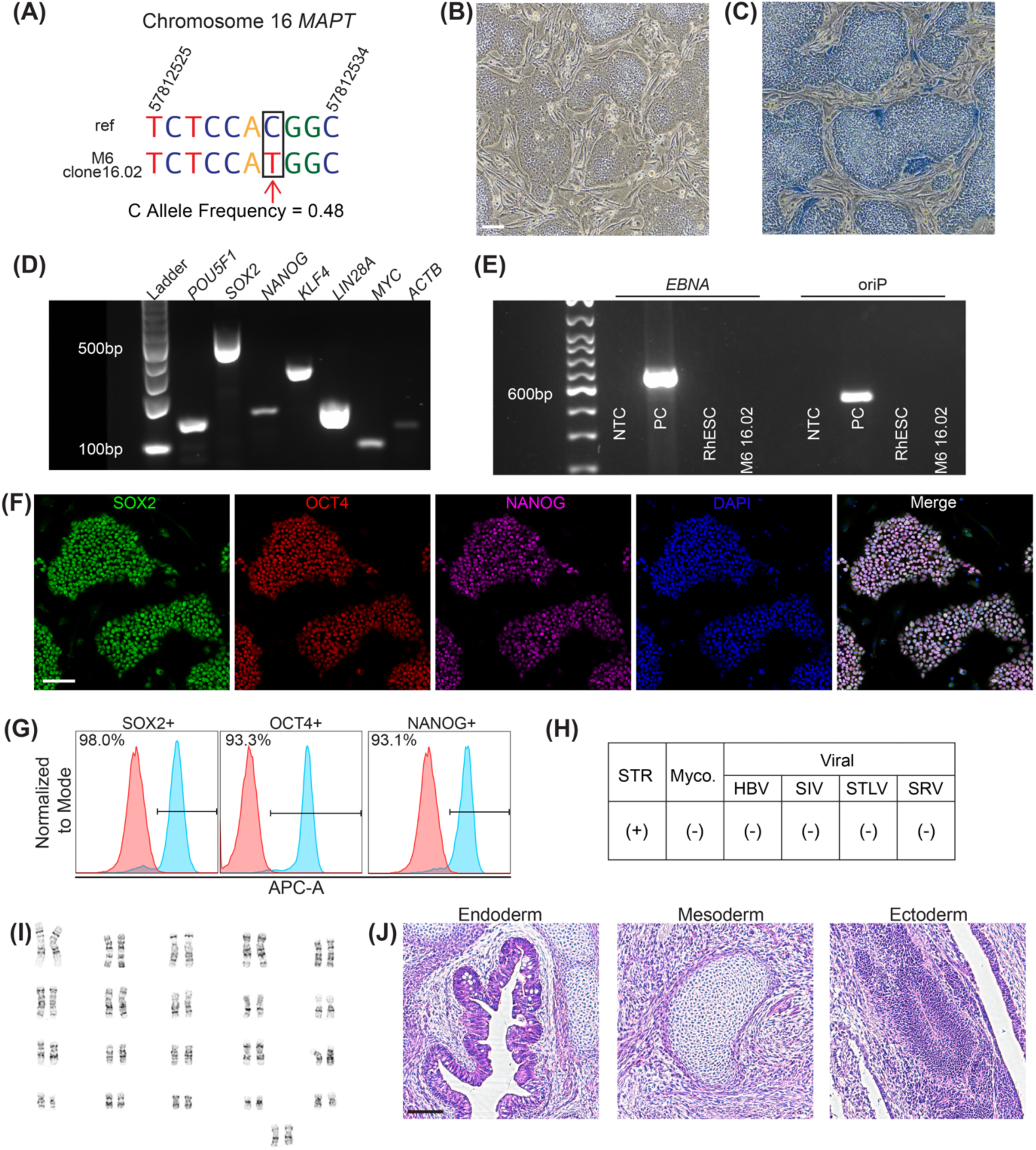
Characterization of RhiPSC line M6 clone 16.02. (A) Amplicon sequencing of *MAPT* R406W with mutant T allele indicated in the black box with a red arrow compared to reference genome (ref). (B) Phase contrast image of colonies grown on MEFs and (C) positive alkaline phosphatase staining; scale bar, 100 µm. (D) RT-PCR detection of endogenous pluripotent mRNAs *POU5F1*, *SOX2*, *NANOG*, *KLF4*, *LIN28*, and *MYC*. *ACTB* was used as a reference gene. (E) Amplification of EBNA and oriP in RhiPSCs at P18, no template control (NTC), and the rhesus embryonic stem cell line r420 (RhESC). Plasmid EM2K was used as a positive control (PC) for the detection of EBNA and oriP. (F) Positive immunofluorescence staining for pluripotency proteins SOX2 (green), OCT4 (red), and NANOG (far red), colocalized with DAPI (blue). Scale bar, 100 µm. (G) Flow cytometry-detected APC-A+ cells were 93.1% positive for SOX2, 98% positive for OCT4, and 93.3% positive for NANOG. (H) Short tandem repeat (STR) analysis confirmed (+) RhiPSC line identity. Mycoplasma (Myco.), Herpes B Virus (HBV), Simian Immunodeficiency Virus (SIV), Simian T-Lymphotropic Virus (STLV), and Simian Retrovirus (SRV) were not detected (-). (I) Normal 42,XX karyotype at P15. (J) H&E staining of a teratoma containing endoderm, mesoderm, and ectoderm. Scale bar, 100 µm.

RhiPSCs retained their respective *MAPT* WT or mutant alleles following episomal reprogramming (**Figures 4A, 5A; Supplementary Figures 4A-9A**). Colonies typically presented as homogeneous, compact PSC-like colonies with a high nucleus-to-cytoplasm ratio (**Figures 4B, 5B; Supplementary Figures 4B-9B**). Individual RhiPSC colonies were alkaline phosphatase positive with distinct colony boundaries (**Figures 4C, 5C; Supplementary Figures 4C-9C**). RT-PCR confirmed robust expression of pluripotency genes *POU5F1*, *NANOG*, *SOX2*, *KLF4*, *MYC* and *LIN28A* (**Figures 4D, 5D; Supplementary Figures 4D-9D**). In addition, the episomal plasmids were no longer detected in the established clonal RhiPSC lines when tested between P14-P18 (**Figures 4E, 5E; Supplementary Figures 4E-9E**). Immunostaining and flow cytometric analysis detected uniformly expressed SOX2, OCT4, and NANOG (**Figures 4F,G, 5F,G; Supplementary Figures 4F,G-9F,G**). The cells were positively authenticated and were negative to mycoplasma, HBV, SIV, STLV and SRV (**Figures 4H, 5H; Supplementary Figures 5H-9H**). Both female (XX) and male (XY) RhiPSC lines had normal karyotypes (**Figures 4I, 5I; Supplementary Figures 4I-9I**). Established RhiPSCs could be maintained without indication of differentiation for more than 30 passages over 4 months of continued cell culture when grown on MEFs in UPPS medium. To test the capacity of RhiPSCs to form teratomas, RhiPSCs were injected into NCG-X mice. All established lines formed tumors within 8 weeks post-injection. Histological analysis confirmed that all the harvested tumors contained multilineage derivatives of all three germ layers including endoderm, ectoderm, and mesoderm lineages (**Figures 4J, 5J; Supplementary Figure 4J-9J**). Together, these observations verify pluripotency and the ability to form derivatives from all primary germ layers.

### 3.5 RhNPC Derivation and Characterization

RhiPSCs derived from macaques W4, M4, W6, and M6 were differentiated to NPCs over 21 days. RhNPCs mainly presented with uni- or bipolar neurites extending from a small soma. All four RhNPCs lines expressed ectodermal lineage transcription factor PAX6 in the nuclei and intermediate filament protein NESTIN in the cytosol (**Figure 6**), whereas the expression of the pluripotency-associated transcription factor OCT4 was not detected at 21 days of differentiation (not shown).

**Figure 6.**
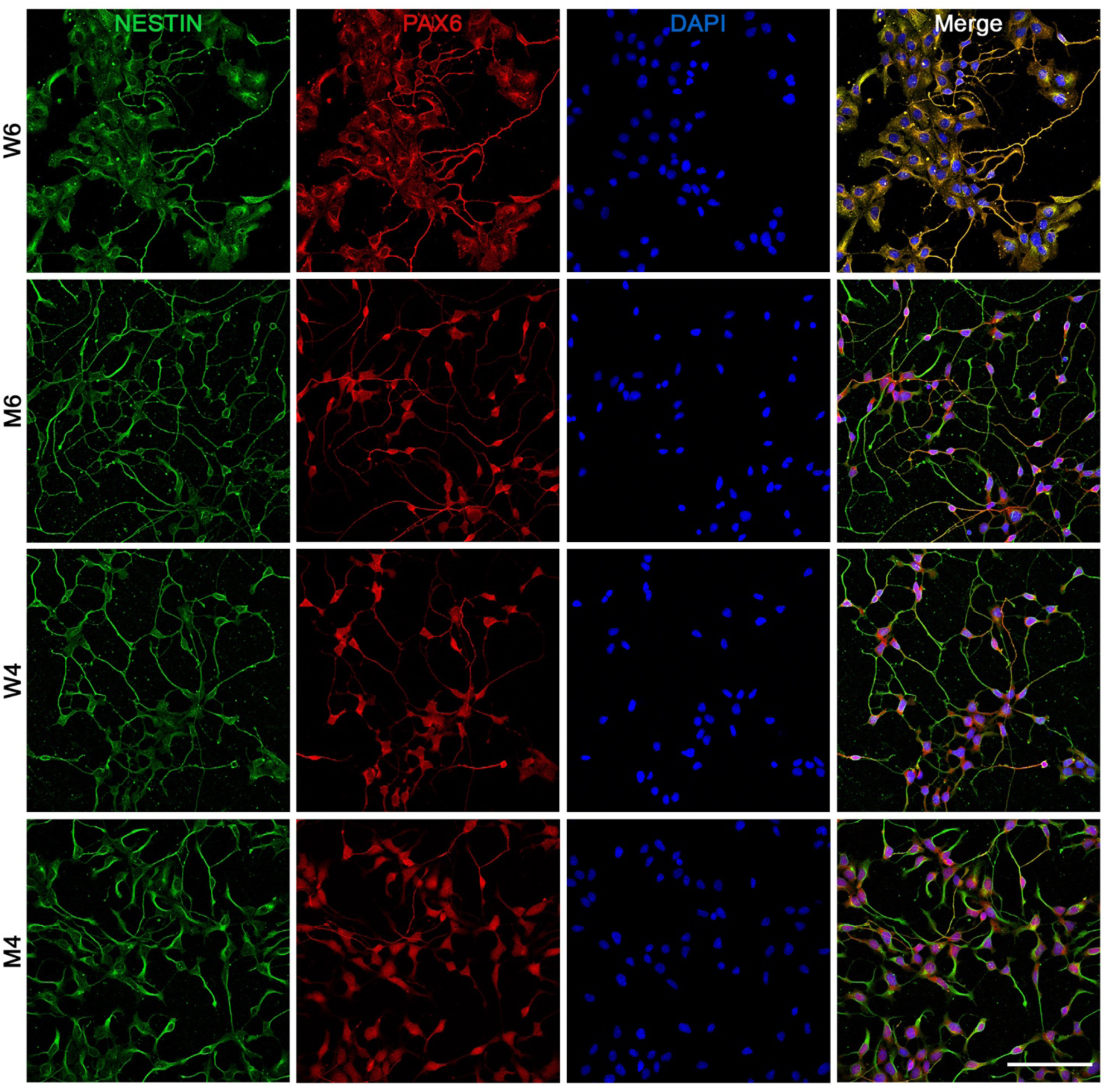
Neural progenitor cells derived from RhiPSC lines. Immunofluorescence images showing positive NESTIN (green) and PAX6 (red) expression with DAPI (blue) after 21 days of differentiation. Representative images from W6 clone 5.07, M6 clone 16.02, W4 clone 14.06, and M4 clone 14.09. Scale bar, 100 µm.

## 4.0 Discussion

The present study identified a replicable workflow for generating and maintaining rhesus fibroblast-derived RhiPSCS to support the *in vitro* modeling of *MAPT* R406W FTD. Our stepwise approach optimized methods for fibroblast derivation, fibroblast reprogramming, and RhiPSC maintenance. Using non-integrating episomal (oriP/EBNA1) plasmids, we generated eight RhiPSC lines that fulfill ISSCR standards, including four *MAPT* R406W lines. These RhiPSCs were patterned to RhNPCs to demonstrate their differentiation potential and enable future studies on mechanisms of tau-related neuropathology in genetic FTD. Importantly, this *in vitro* model will be critical for validating therapeutic targets before *in vivo* evaluation, thereby reducing the number of valuable rhesus monkeys used in preclinical studies aimed at human clinical trials. In this discussion, we will address how our methods compare to previous publications reporting successful RhiPSC generation, relate this to methodology for generating and maintaining human iPSCs, and consider implications for FTD disease modeling.

Macaque iPSC generation is generally less efficient than hiPSC generation (Stauske et al., 2020), probably due to the limited number of labs working on these species, which in turn results in less development of optimized protocols for stably maintaining undifferentiated cells. Four studies have reported the generation of RhiPSCs using Sendai viruses (Jocher et al., 2024; Lara et al., 2023; Tao et al., 2021; Zhao et al., 2018), and four studies have reported the generation of RhiPSCs via episomal reprogramming with pCXLE plasmids or piggyBac transposons (Rodriguez-Polo et al., 2021; Rodríguez-Polo et al., 2022; Stauske et al., 2020; Zhang et al., 2017).

Several factors may have contributed to our inefficient generation of stable RhiPSCs via Sendai reprogramming, in contrast to previous reports (Jocher et al., 2024; Lara et al., 2023; Tao et al., 2021; Zhao et al., 2018). First, as Sendai viral vector production depends on commercial vendors, differences between batches may produce divergent outcomes across labs. Second, unreported variations in cell culture technique (e.g., criteria for cell passaging following viral transduction) may affect cell viability. Third, previous publications did not mention if the generated RhiPSCs were cryopreserved and stably thawed, a criterion listed by the ISSCR standards. Knowledge of successful cryopreservation and thawing methods may have prevented the negative outcome of our Sendai trial 3. Lastly, it should be acknowledged that negative results are seldom reported (Emborg, 2023). Most studies describe clonal lines derived from one animal (Jocher et al., 2024; Lara et al., 2023; Zhao et al., 2018); whether more than one trial or animal/cell donor were needed to successfully generate RhiPSC clones was not stated. Similarly, excessive proliferation following fibroblast transduction, as observed in our first two Sendai trials, was not previously reported. This was intriguing, as the *MAPT* R406W fibroblasts, which could not be maintained past P3-P4, were maintained up to P8. The fibroblasts may have been immortalized by the Sendai system, due to the expression of the oncogene c-Myc (Wang et al., 2011). When inducing pluripotency, a precise balance of c-Myc and Klf4 expression is required, as Klf4 has antiproliferative effects (Takahashi & Yamanaka, 2006). Despite our efforts to balance the expression of c-Myc and Klf4, excessive proliferation was observed following transduction.

Historically, non-integrating Epstein-Barr virus-based vectors encoding an origin of replication (oriP) and the Epstein-Barr nuclear antigen 1 (EBNA1) protein (i.e., oriP/EBNA1 plasmids) have been used for human fibroblast reprogramming (Howden et al., 2006; Yu et al., 2009). In humans and monkeys, EBNA1 binding to the oriP tethers the plasmid to mitotic chromosomes without integration. In the absence of selective pressure, the episomes are gradually lost from the cells at a rate of ∼5% per cell division due to defects in plasmid synthesis and partitioning, leaving behind iPSCs free of plasmid and transgene sequences (Yu et al., 2009). For plasmid-based reprogramming, several variables must be considered that affect success: reprogramming factors and promoter, electroporation conditions, and transfected cell plating density.

Selection of reprogramming factors and promoter plays a critical role in RhiPSC reprogramming success. Although the four Yamanaka factors are usually considered sufficient for generating iPSCs, monkey cells often require additional factors to improve reprogramming efficiency (Debowski et al., 2015; Takahashi & Yamanaka, 2006). Macaque iPSCs have been successfully derived using pCXLE plasmids to express five reprogramming factors under a CAG promoter, including Klf4, Sox2, Oct4, L-Myc, and Lin28 (Rodríguez-Polo et al., 2022; Stauske et al., 2020; Zhang et al., 2017). To increase reprogramming efficiency, these studies replaced C-Myc with non-transforming L-Myc, and included a p53 suppression element (shp53) (Okita et al., 2011). In our episomal trials 1-6, we used oriP/EBNA1 plasmids expressing Klf4, Sox2, Oct4, C-Myc, and Lin28 under an EF1α promoter, and added Nanog in trials 7-13. Nanog, although not required for the appearance of iPSC colonies, has been shown to improve reprogramming efficiency (Takahashi & Yamanaka, 2006; Yu et al., 2007). Furthermore, we chose EF1α instead of a CAG promoter as it is more resistant to methylation-mediated transcriptional silencing leading to more stable expression of reprogramming factors (Sun et al., 2021). Our addition of the plasmid encoding a miR-302/367 cluster enhanced reprogramming efficiency through indirect positive regulation of Oct4 (Howden et al., 2015; Hu et al., 2013).

Neon programs #5, #14, #15, #16, #20, #23, and #24 achieved successful plasmid uptake across several rhesus fibroblast lines. These parameters can be the basis for future reprogramming trials, although optimization of electroporation conditions per cell line is recommended to tailor an approach specific to the fibroblast line. In that regard, plating densities need to account for cell loss following electroporation. Further, due to the stochastic nature of fibroblast reprogramming, the outcome of different trials with similar parameters may vary, thus multiple replicates are also recommended. Some fibroblast lines seem to be more resistant to reprogramming than others. For example, trials 4-6 failed to reprogram fibroblast line M2 and warrants further analyses regarding the subject-dependent impact of the *MAPT* R406W mutation. It should be noted that previous studies have utilized Lonza Nucleofector devices for plasmid electroporation (Lonza, Basel, Switzerland; (Rodriguez-Polo et al., 2021; Rodríguez-Polo et al., 2022; Stauske et al., 2020; Zhang et al., 2017). Lonza devices contain electroporation programs in which the electroporation parameters (e.g., pulse voltage) are proprietary information; thus, replicability could be compromised if the company changes the parameters. In contrast, the Neon Transfection system used in this study has 24 different pre-set programs with the parameters publicly available (see **Supplementary Table 2**). It should be noted that the Neon NxT system (ThermoFisher Scientific #NEON18SK) that replaced the Neon Transfection System has the same 24 different pre-set programs.

Culture conditions for RhiPSC maintenance have not been standardized, unlike hiPSC culture. We hypothesized that our initial RhiPSC instability was due to the E12+IWR1 medium used for maintenance. E12+IWR1 medium is based on Essential 8 (E8) medium, which was developed as a fully defined media for the culture of human pluripotent stem cells (PSCs) (Chen et al., 2011). However, E8 is not optimal for NHP PSC maintenance (Stauske et al., 2020). By supplementing E8 with Nodal, glutathione, chemically defined lipids, and Glutamax (E12), plus the WNT inhibitor IWR1, the culture of cynomolgus macaque iPSCs was improved (Baik et al., 2023). Yet, E12+IWR1 medium did not support the long-term culture of RhiPSCs in this study. Stauske et al. developed a ‘Universal Primate Pluripotent Stem Cell (UPPS)’ medium (Stauske et al., 2020), consisting of StemMACS iPS Brew XF medium supplemented with the Wnt signaling inhibitor IWR-1 and the GSK3β inhibitor CHIR99021. This medium was capable of long-term maintenance of RhiPSCs, RhESCs, and baboon iPSCs in an undifferentiated state (Rodriguez-Polo et al., 2021; Rodríguez-Polo et al., 2022; Stauske et al., 2020). Thus, we switched to UPPS medium for RhiPSCs generated during episomal reprogramming trial 7 and later, and the eight RhiPSC lines demonstrated typical PSC morphology over several passages and normal karyotypes. A limitation of the UPPS medium is that the composition of the StemMACs iPSC Brew XF is proprietary. Future research should focus on identifying components critical to maintaining RhiPSCs to develop a fully defined media.

Maintaining undifferentiated RhiPSCs seems to depend on the balance between WNT inhibition and activation. In the canonical Wnt/β-catenin pathway, Wnt ligand binds to its receptor complex, which prevents phosphorylation of glycogen synthase kinase (GSK3). In the absence of phosphorylated GSK3, β-catenin is free to translocate to the nucleus where it influences gene expression. Otherwise, phosphorylated GSK3 leads to proteasome-mediated degradation of β-catenin. The presence of both IWR1 and CHIR99021 in the UPPS medium raises questions, as IWR1 is a WNT inhibitor and CHIR99021 is a GSK3β inhibitor. The combination of IWR1 and CHIR99021 has been shown to be highly effective in promoting self-renewal of human ESCs and mouse epiblast cells through a poorly understood mechanism (Kim et al., 2013). PSC media supplemented with both IWR1 and CHIR99021 may be applicable to several nonhuman PSC species, as this combination has also been reported for the maintenance of undifferentiated pig iPSCs (Conrad et al., 2023; Edwards et al., 2025). Thus, the mechanism by which Wnt signaling influences the pluripotent state across species requires further investigation.

Cloning efficiency experiments demonstrated that MEFs best support RhiPSC self-renewal compared to Matrigel or Vitronectin. Unlike human PSC culture, which can be maintained under feeder-free conditions with ease, most publications on NHP iPSCs report the use of MEFs or MEF-conditioned media for cell derivation and/or maintenance (Aron Badin et al., 2019; Hong et al., 2014; Jacobo Lopez et al., 2022; Jocher et al., 2024; Nakai et al., 2018; Navara et al., 2018; Rodriguez-Polo et al., 2021; Tao et al., 2021; Zhang et al., 2017). Establishing a feeder-free system for the culture of NHP PSCs would reduce costs, prevent inconsistencies due to batch variations, and eliminate MEF background in experimental analyses and cell differentiations. Matrigel and Vitronectin were evaluated as these matrices are feeder-free options that support hiPSC self-renewal. Our results demonstrate that the addition of Y-27632 or CEPT to cell culture media at the time of passage increases RhiPSC self-renewal on Matrigel and Vitronectin. Experiments building on these findings with evaluation of genetic stability and stemness of colonies will pave the way for identifying feeder-free culture conditions for monkey iPSCs.

The RhiPSCs generated from *MAPT* R406W monkeys are envisioned as a resource for studies of FTD linked to tau pathology across species and for testing novel therapeutics before preclinical testing in the valuable animals. As such, we patterned PAX6 and NESTIN-expressing rhesus neural progenitor cells (RhNPCs) over 21 days. Our results indicate that fibroblasts derived from *MAPT* R406W monkeys can also be useful for unraveling tau-related pathogenesis. The observation of enlarged and multinucleated *MAPT* R406W fibroblasts that ceased proliferation around P3-P4 suggests failed mitosis. In fibroblasts (Connolly et al., 1977; Sjöberg et al., 2006), tau is localized to interphasic cytoskeleton (Rossi et al., 2008) and bound to nuclear DNA (Wei et al., 2008). During fibroblast pro/metaphase, tau is found in centrosomes, the mitotic spindle, and the periphery of condensed chromosomes (Rossi et al., 2008). Interestingly, aneuploidy has been observed in cell lines with *MAPT* mutations (Rossi et al., 2014). Further interrogation of fibroblasts from mutation carriers may help understand the nuclear mechanisms contributing to the observed phenotype and its impact on neuronal pathology.

The ISSCR reporting standards for publishing with iPSCs (Ludwig et al., 2023) aims to ensure safe sharing of cell resources and credibility of data. To the best of our knowledge, only one report besides our own has characterized an RhiPSC line following the ISSCR standards (Lara et al., 2023). The application of ISSCR standards to NHP iPSCs can be extremely useful, given that NHP iPSC culture is not as standardized as hiPSC culture and these reporting practices would convey key information for replicable results. Details that seem minor, such as cryopreservation methods or split ratios, would contribute to a lab’s success in reproducing a given study. Special considerations should be taken when reporting on RhiPSCs. For example, one of the ISSCR reporting requirements is sterility, which includes viral screening of cell cultures. RhiPSCs have the potential to carry zoonotic agents such as Herpes B virus that are not found in hiPSC cultures; thus, this virus should also be tested for and reported upon. This becomes especially important if the RhiPSCs are to be shared as a resource among researchers.

## 5.0 Conclusions

Overall, these results demonstrated the replicability of oriP/EBNA1 episomal reprogramming for the generation of RhiPSCs. Variables affecting fibroblast reprogramming trials included selection of reprogramming factors and promoter, electroporation conditions, and transfected fibroblast plating density. Medium (UPPS) and matrix (MEFs) were identified for continued culture of undifferentiated and genetically stable RhiPSCs. All eight RhiPSCs lines (*MAPT* WT n=4, R406W n=4) were fully characterized according to ISSCR standards, and one clonal line per animal was differentiated to RhNPCs over 21 days to support the *in vitro* modeling of *MAPT* R406W FTD. These cells will be a critical resource for understanding how the *MAPT* R406W mutation drives disease, identifying biomarkers, and testing novel therapies before preclinical testing in the valuable monkeys.

## 6.0 Resource Availability Statement

All eight RhiPSC lines (W4 clone #14.06; W4 clone #16.10; M4 clone #14.09; M4 clone #16.01; W6 clone #5.07; W6 clone #15.03; M6 clone #16.02; M6 clone #4.04) are available to investigators upon reasonable request through the Cell Line Repository at the Wisconsin National Primate Research Center.

## Acknowledgements

Reprogramming plasmids were kind gifts from Drs. J. Thomson, S. Howden, and J. Vadolas. The authors gratefully acknowledge the University of Wisconsin Waisman Center Stem Cell Core for performing the Sendai virus reprograming trial #3.

