## Supplementary Tables and Figures for "Replicable generation of rhesus macaque iPSCs for *in vitro* modeling of genetic frontotemporal dementia"

**Supplementary Table 1**. Media formulations.

| Medium | Reagent | Concentration | Vendor | Cat. No. |
|---|---|---|---|---|
| Biopsy | DMEM/F12 | 99% | Thermo Fisher Scientific | SH30023 |
| | Pen-Strep | 1% | Thermo Fisher Scientific | 15140122 |
| Digestion | hi-glucose DMEM | 80% | Thermo Fisher Scientific | 11960044 |
| | FBS | 20% | Peak Serum | PSFB4 |
| | Collagenase type I | 0.4 mg/mL | Worthington Biochemical | LS004194 |
| | DNAse I | 2 mg/mL | Sigma-Aldrich | DN25 |
| | Pen-Strep | 1% | Thermo Fisher Scientific | 15140122 |
| Fibroblast | DMEM/F12 | 80% | Thermo Fisher Scientific | SH30023 |
| | FBS | 20% | Peak Serum | PSFB4 |
| | Sodium Pyruvate | 1% | Lonza Bioscience | BEBP13-115E |
| | Non-essential amino acids | 1% | Gibco | 11140050 |
| | FGF-2 | 20 ng/mL | Bio-Techne | BT-FGFB-100 |
| | Pen-Strep | 1% | Thermo Fisher Scientific | 15140122 |
| N2B27 | DMEM/F12 | 50% | Thermo Fisher Scientific | SH30023 |
| | Neurobasal | 50% | Thermo Fisher Scientific | 21103049 |
| | B27 Supplement | 1X | Thermo Fisher Scientific | 12587010 |
| | N2 Supplement | 1X | Thermo Fisher Scientific | 17502048 |
| | GlutaMAX | 2 mM | Thermo Fisher Scientific | 35050061 |
| | Non-essential amino acids | 0.1 mM | Gibco | 11140050 |
| | BME | 0.1 mM | Thermo Fisher Scientific | 21985023 |
| | BSA | 2 mg/mL | Sigma-Aldrich | A8806 |
| | FGF-2 | 20 ng/mL | Bio-Techne | BT-FGFB-100 |
| | IWR1 | 2.5 μM | STEMCELL Technologies | 72562 |
| | CHIR99021 | 3 μM | Tocris | 4423 |
| UPPS | StemMACS iPS Brew XF | base | Miltenyi Biotec | 130-104-368 |
| | endo-IWR1 | 1 μM | STEMCELL Technologies | 72562 |
| | CHIR99021 | 0.5 μM | Tocris | 4423 |
| E11+Bu | DF3S | base | Thermo Fisher Scientific | custom order |
| | Transferrin | 10.7 μg/mL | Thermo Fisher Scientific | 2914-HT-001G |
| | Insulin | 20 μg/mL | Sigma-Aldrich | I9287-5ML |
| | FGF-2 | 100 ng/mL | Bio-Techne | BT-FGFB-100 |
| | GlutaMAX | 1X | Thermo Fisher Scientific | 35050061 |
| | Chemically Defined Lipids | 1X | Thermo Fisher Scientific | 11905031 |
| | rhNodal | 50 ng/mL | R&D Systems | 3218-ND |
| | Glutathione | 1.94 μg/mL | Sigma-Aldrich | G4251 |
| | Sodium Butyrate | 100 μM | STEMCELL Technologies | 72242 |
| E12+IWR1 | DF3S | base | Thermo Fisher Scientific | custom order |
| | Transferrin | 10.7 μg/mL | Thermo Fisher Scientific | 2914-HT-001G |
| | Insulin | 20 μg/mL | Sigma-Aldrich | I9287-5ML |
| | FGF-2 | 100 ng/mL | Bio-Techne | BT-FGFB-100 |
| | GlutaMAX | 1X | Thermo Fisher Scientific | 35050061 |
| | Chemically Defined Lipids | 1X | Thermo Fisher Scientific | 11905031 |
| | rhNodal | 50 ng/mL | R&D Systems | 3218-ND |
| | Glutathione | 1.94 μg/mL | Sigma-Aldrich | G4251 |
| | TGFβ | 1.7 ng/mL | R&D Systems | 240-B |
| | IWR1 | 2.5 μM | STEMCELL Technologies | 72562 |
| Complete | DMEM/F12 | 90% | Thermo Fisher Scientific | SH30023 |
| | FBS | 10% | Peak Serum | PSFB4 |
| Neural Induction | STEMdiff Neural Induction Medium | base | STEMCELL Technologies | 08581 |
| | STEMdiff SMADi Neural Induction Supplement | 0.5 mL | STEMCELL Technologies | 08581 |
| Neural Freezing | DMSO | 20% | Sigma-Aldrich | D8418 |
| | KnockOut Serum Replacement | 80% | Thermo Fisher Scientific | 10828010 |

**Supplementary Table 2**. Electroporation parameters and outcomes. The percentage of live cells (%Live) and eGFP-expressing cells (%eGFP) post-electroporation are shown for each experimental replicate (rep 1, rep 2), fibroblast line (W4, M4, W6, M6), and Neon transfection program. Data were visually inspected and cutoffs were rendered to reflect programs with the highest %eGFP and relatively high %Live. The Neon programs with both criteria above the cutoff were identified (highlighted in yellow). Asterisks (*) represent replicates in which the %eGFP was above the cutoff. Abbreviations: eGFP, enhanced green fluorescent protein; no., number of pulses; rep, replicate; v, voltage.

| | Program Conditions | W4 | | | | M4 | | | | W6 | | | | M6 | | | |
|---|---|---|---|---|---|---|---|---|---|---|---|---|---|---|---|---|---|
| Program # | v, width, no. | %Live, rep 1 | %eGFP, rep 1 | %Live, rep 2 | %eGFP, rep 2 | %Live, rep 1 | %eGFP, rep 1 | %Live, rep 2 | %eGFP, rep 2 | %Live, rep 1 | %eGFP, rep 1 | %Live, rep 2 | %eGFP, rep 2 | %Live, rep 1 | %eGFP, rep 1 | %Live, rep 2 | %eGFP, rep 2 |
| 1 | none | 97.4 | 0.5 | 91.5 | 0.49 | 97.9 | 0.51 | 87.9 | 0.6 | 99.6 | 0.48 | 99.7 | 0.50 | 96.2 | 0.52 | 98.9 | 0.51 |
| 2 | 1400, 20, 1 | 71.9 | 0.8 | 86.70 | 22.70 | 90.3 | 3.25 | 73.9 | 1.04 | 99.7 | 7.32 | 92.7 | 8.74 | 93.1 | 18 | 98.4 | 4.67 |
| 3 | 1500, 20, 1 | 52 | 2.4 | 77.90 | 22.90 | 88.3 | 3.72 | 75.4 | 3.53 | 98.7 | 15.6 | 90.5 | 13.60 | 90.7 | 26.6 | 97.8 | 12.5 |
| 4 | 1600, 20, 1 | 49.4 | 9.45 | 88.40 | 0.65 | 83.8 | 4.78 | 61.8 | 2.13 | 98.1 | 17.7 | 89.1 | 10.90 | 83.8 | 39.2 | 97.3 | 10 |
| 5 | 1700, 20, 1 | 51.6 | 14.4* | 85.90 | 1.79 | 75.7 | 6.06* | 75.1 | 9.84* | 93.7 | 34.4 | 84.4 | 25.50 | 75.4 | 52.9* | 94 | 20.4* |
| 6 | 1100, 30, 1 | 52.8 | 0.89 | 80.3 | 23.10 | 93.9 | 1.97 | 77.6 | 1.14 | 99.7 | 4.35 | 92.3 | 4.84 | 96 | 10.3 | 96.8 | 1.46 |
| 7 | 1200, 30, 1 | 81.5 | 2.44 | 83.9 | 4.17 | 92.3 | 2.67 | 75.1 | 1.79 | 99.5 | 7.23 | 92.6 | 10.90 | 92.4 | 24 | 95.1 | 3.84 |
| 8 | 1300, 30, 1 | 68.8 | 1.44 | 76.00 | 22.00 | 88.5 | 3.68 | 68.2 | 3.86 | 99.2 | 10.3 | 90.3 | 19.40 | 87 | 32.2 | 96 | 3.54 |
| 9 | 1400, 30, 1 | 59.5 | 9.09 | 74.10 | 27.90 | 79.2 | 4.89 | 59.9 | 0.12 | 95.3 | 28.6 | 91.8 | 21.80 | 72.9 | 48.8 | 87.5 | 20.2* |
| 10 | 1000, 40, 1 | 75.7 | 1.53 | 80.10 | 21.40 | 94 | 1.85 | 75.7 | 0.9 | 99.6 | 6.03 | 90.0 | 4.87 | 94.8 | 11.4 | 95.4 | 2.18 |
| 11 | 1100, 40, 1 | 64.6 | 2.35 | 75.70 | 27.90 | 91.1 | 2.44 | 75.4 | 0.48 | 99 | 12 | 99.4 | 3.78 | 91 | 22.7 | 95.7 | 6.37 |
| 12 | 1200, 40, 1 | 54.1 | 2.54 | 61.40 | 28.90 | 87.3 | 3.48 | 78.8 | 6.39 | 97.9 | 17.5 | 89.9 | 27.00 | 79.7 | 39.3 | 94.5 | 2.82 |
| 13 | 1100, 20, 2 | 72.9 | 2.56 | 85.30 | 5.00 | 92.1 | 2.29 | 79.8 | 2.84 | 98.8 | 15.6 | 93.4 | 9.60 | 91.9 | 20.2 | 95.9 | 4.22 |
| 14 | 1200, 20, 2 | 45 | 4.61 | 74.4 | 31.2* | 85.9 | 3.73 | 76.5 | 4.98 | 96.2 | 24.7 | 91.1 | 20.90 | 84.2 | 33.9 | 94.6 | 9.24 |
| 15 | 1300, 20, 2 | 69 | 8.92 | 66.9 | 27.5 | 74.1 | 5.67* | 77.8 | 11.4* | 86.5 | 48.6* | 83.3 | 35.40* | 70 | 49.3 | 93.7 | 19.1 |
| 16 | 1400, 20, 2 | 52.6 | 10.8* | 58.8 | 33.7* | 60.5 | 7.34* | 79.6 | 0.42 | 70.7 | 63.1* | 87.5 | 30.60 | 87.5 | 58.3* | 87.9 | 21.9* |
| 17 | 850, 30, 2 | 83.9 | 0.49 | 81.2 | 14.4 | 94.9 | 1.64 | 76.6 | 0.48 | 99.8 | 4.23 | 91.4 | 2.35 | 96.3 | 6.63 | 96 | 1.01 |
| 18 | 950, 30, 2 | 75.6 | 1.25 | 77.80 | 30.5* | 97.8 | 0.44 | 76.3 | 1.23 | 97.6 | 19.4 | 89.1 | 7.24 | 93.1 | 17.2 | 95.4 | 2.75 |
| 19 | 1050, 30, 2 | 75.1 | 4.92 | 83.20 | 2.00 | 88.1 | 2.58 | 78 | 0.06 | 95.8 | 23 | 89.1 | 20.80 | 84.2 | 32.4 | 94.5 | 9.49 |
| 20 | 1150, 30, 2 | 46.2 | 6.73 | 34.1 | 34.9* | 73 | 5.14 | 71.9 | 0.6 | 83.2 | 47.5 | 81.8 | 40.30* | 88.5 | 50.7 | 96.8 | 0.75 |
| 21 | 1300, 10, 3 | 59.6 | 0.62 | 77.8 | 20.3 | 92.9 | 2.14 | 70 | 3.46 | 98.8 | 14.5 | 90.7 | 4.79 | 94.2 | 13.4 | 95 | 5.86 |
| 22 | 1400, 10, 3 | 63.9 | 2.9 | 66.9 | 29.1 | 88.2 | 3.09 | 76.4 | 5.64 | 95.1 | 30.2 | 89.8 | 18.70 | 86.3 | 28.8 | 95.5 | 0.9 |
| 23 | 1500, 10, 3 | 44.6 | 0 | 73.2 | 11.2 | 82.9 | 4.3 | 72.9 | 9.49* | 92.5 | 36 | 83.0 | 31.20 | 81.1 | 39.9 | 94.2 | 14.5 |
| 24 | 1600, 10, 3 | 53.4 | 19.1* | 57.9 | 29.4 | 74.1 | 4.91 | 68.4 | 0.81 | 84.5 | 51.2* | 78.7 | 47.80* | 77.6 | 50.9* | 97 | 0.68 |
| Cutoff | | >50% | >10% | >55% | >30% | >60% | >5.5% | >70% | >9% | >70% | >48% | >78% | >35% | >75% | >50.9% | >85% | >20% |

**Supplementary Table 3**. Primer sequences.

| Amplicon | Forward (5'-3') | Reverse (5'-3') | Amplicon Size (base pairs) |
|---|---|---|---|
| *ACTB* | CTA CCA TGA GCT GCG TGT GG | GTA CAT GGC TGG GGT GTT GA | 130 |
| *EBNA1* | ATC GTC AAA GCT GCA CAC AG | CCC AGG AGT CCC AGT AGT CA | 666 |
| *KLF4* | TCC CGG GGA TTT ATA GCT CG | GTA AGG TTT CTC ACC TGT GTG G | 273 |
| *LIN28A* | ATC AAA AGG AGA CAG GTG CTA C | TCC CGA AAG TAG GTT GGC TT | 180 |
| *MAPT* | GGG CAC AGG GAT TAC GAA TTG | CAC CCC CAG TCT CTT GAA ATC | 1052 |
| *MYC* | GGT AGT GGA AAA CCA GCC TCC | GCA GTA GAA ATA CGG CTG CAC | 103 |
| *NANOG* | GGA TCC AGC TTG TCC CCA AA | AGG AAG GAA GAG GAG AGA CAG T | 161 |
| oriP | TTC CAC GAG GGT AGT GAA CC | TCG GGG GTG TTA GAG ACA AC | 544 |
| *POU5F1* | GCC CGA AAG AGA AAG CGA AC | CAC ACT CGG ACC ACA TCC TTC | 146 |
| *SOX2* | GGT AGG AGC TTT GCA GGA AGT | CCA ACG ATG TCA ACC TGC ATG | 428 |

**Supplementary Table 4**. Antibody information.

| | Primary Antibody | Clonality | Host Species | Dilution | Vendor | Cat. No. | Secondary Antibody | Dilution | Vendor | Cat. No. |
|---|---|---|---|---|---|---|---|---|---|---|
| ICC | OCT3/4 | monoclonal | mouse | 1:1000 | Santa Cruz | sc-5279 | Alexa fluor donkey anti-mouse 594 | 1:1000 | Invitrogen | A-10037 |
| | SOX2 | monoclonal | goat | 1:1000 | Novus Biologicals | AF2018 | Alexa fluor donkey anti-goat 488 | 1:1000 | Invitrogen | A-11055 |
| | NANOG | monoclonal | rabbit | 1:1000 | Cell Signaling Technologies | 4903 | Alexa fluor donkey anti-rabbit 647 | 1:1000 | Invitrogen | A-31573 |
| | PAX6 | polyclonal | rabbit | 1:500 | Biolegend | PRB-278P | Alexa fluor donkey anti-rabbit 594 | 1:500 | Invitrogen | A-21207 |
| | NESTIN | monoclonal | mouse | 1:500 | Thermo Fisher Scientific | MA1-110 | Alexa fluor donkey anti-mouse 488 | 1:500 | Invitrogen | A-21202 |
| Flow Cytometry | OCT3/4 | monoclonal | mouse | 1:100 | Santa Cruz | sc-5279 | APC goat anti-mouse IgG | 1:500 | Biolegend | 405308 |
| | SOX2 | monoclonal | rabbit | 1:100 | Cell Signaling Technologies | 3579 | goat anti-rabbit IgG (H+L) cross absorbed, APC | 1:500 | Thermo Fisher Scientific | A-10931 |
| | NANOG | monoclonal | rabbit | 1:100 | Cell Signaling Technologies | 4903 | goat anti-rabbit IgG (H+L) cross absorbed, APC | 1:500 | Thermo Fisher Scientific | A-10931 |

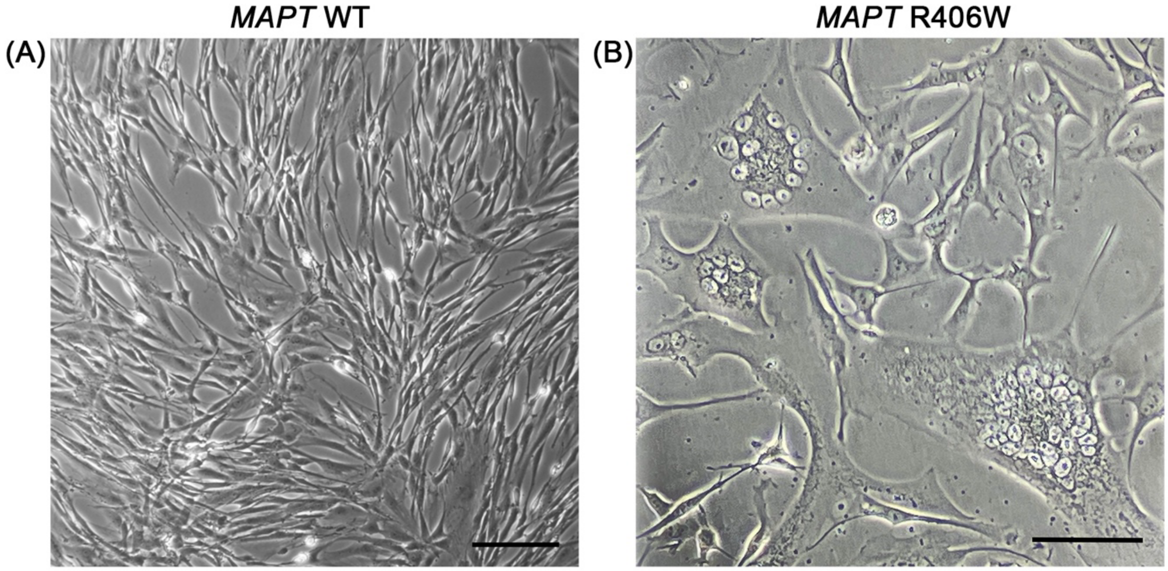


**Supplementary Figure 1.** Representative images of (A) *MAPT* WT fibroblasts at P5 from W4 at P5 and (B) *MAPT* R406W fibroblasts at P3 from M2. Scale bar, 100 µm.

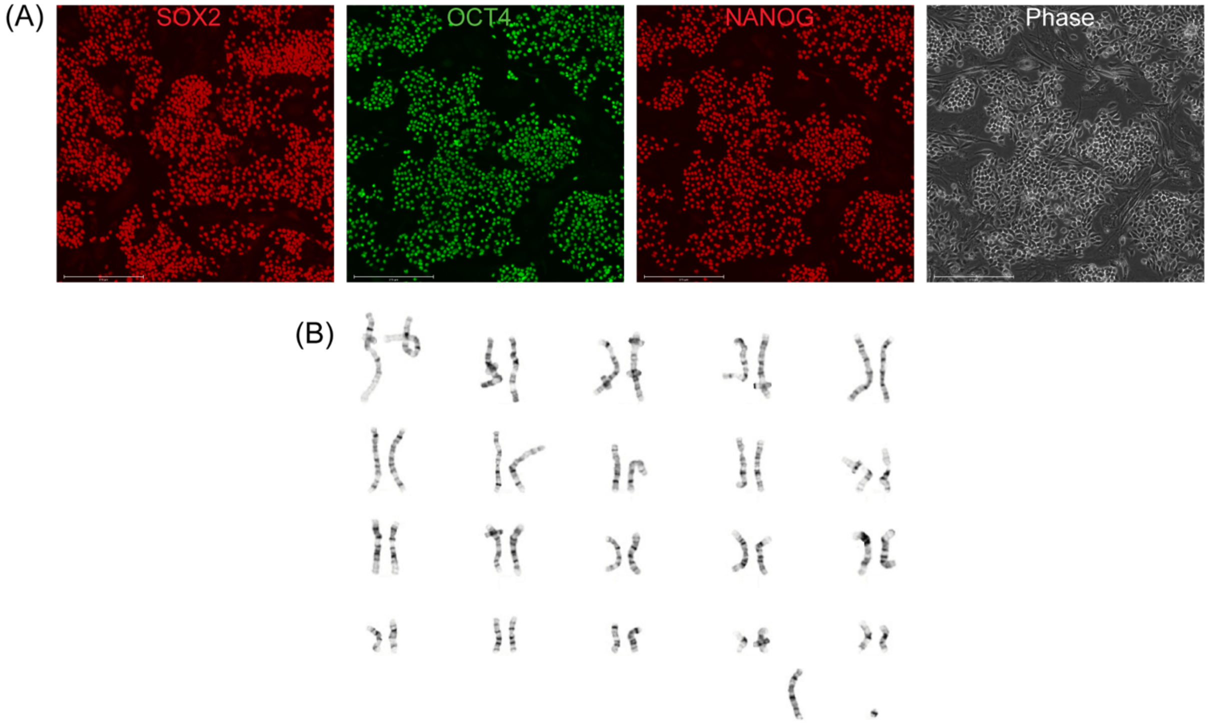


**Supplementary Figure 2.** RhiPSCs derived from animal M4 in episomal reprogramming trial 1 and maintained in E12+IWR1 medium. (A) RhiPSCs expressed pluripotency proteins SOX2, OCT4, and NANOG, yet cells within colonies were not tightly compacted. (B) Abnormal karyotype: 42,XY,der(6)t(3;6)(q;q).

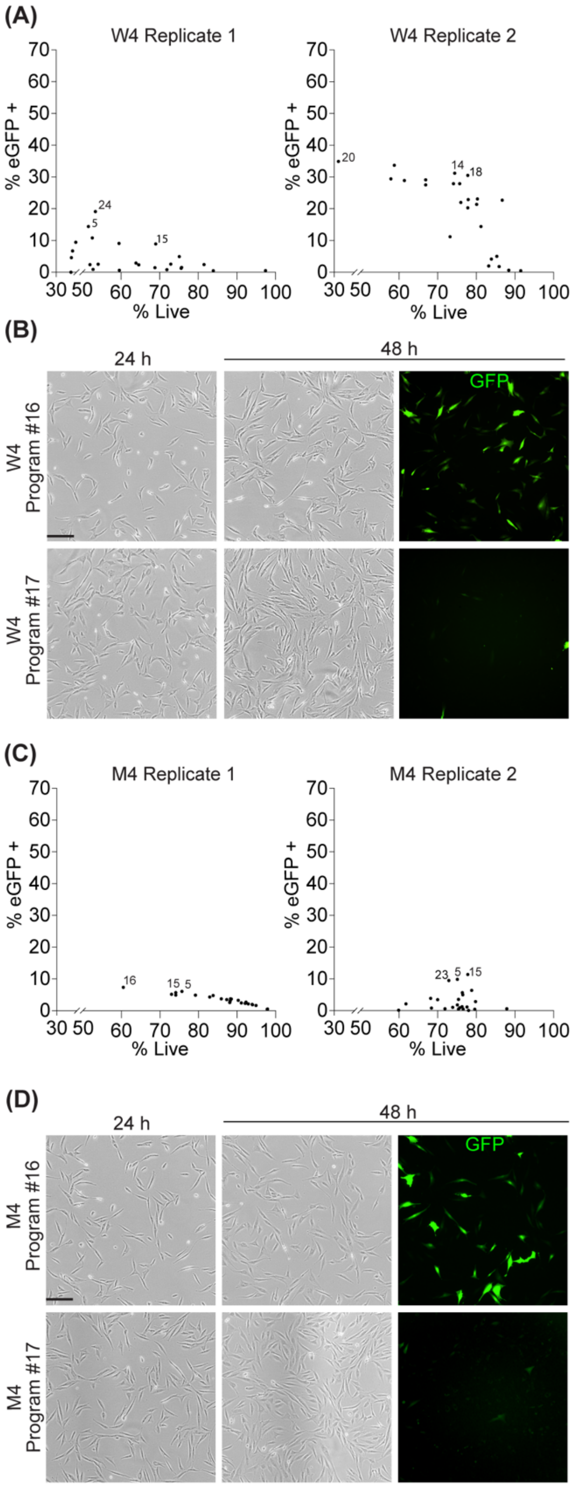


**Supplementary Figure 3.** Optimization of electroporation efficiency for fibroblast lines W4 and M4 using the Neon Transfection system. The percentage of eGFP-expressing cells (% eGFP+; y-axes) were plotted against the percentage of live cells (% live; x-axes); each dot represents data from each of the 24 different Neon programs. (B, D) Representative phase contrast images of fibroblasts lines W4 (B) and M4 (D) demonstrating post-electroporation cell survival at 24 and 48 h on Neon program #16 (an optimal program) and program #17 (a suboptimal program). Corresponding immunofluorescence images were taken 48 h post-electroporation to demonstrate the eGFP+ population resulting from Neon programs #16 and #17. Scale bars, 100 μm.

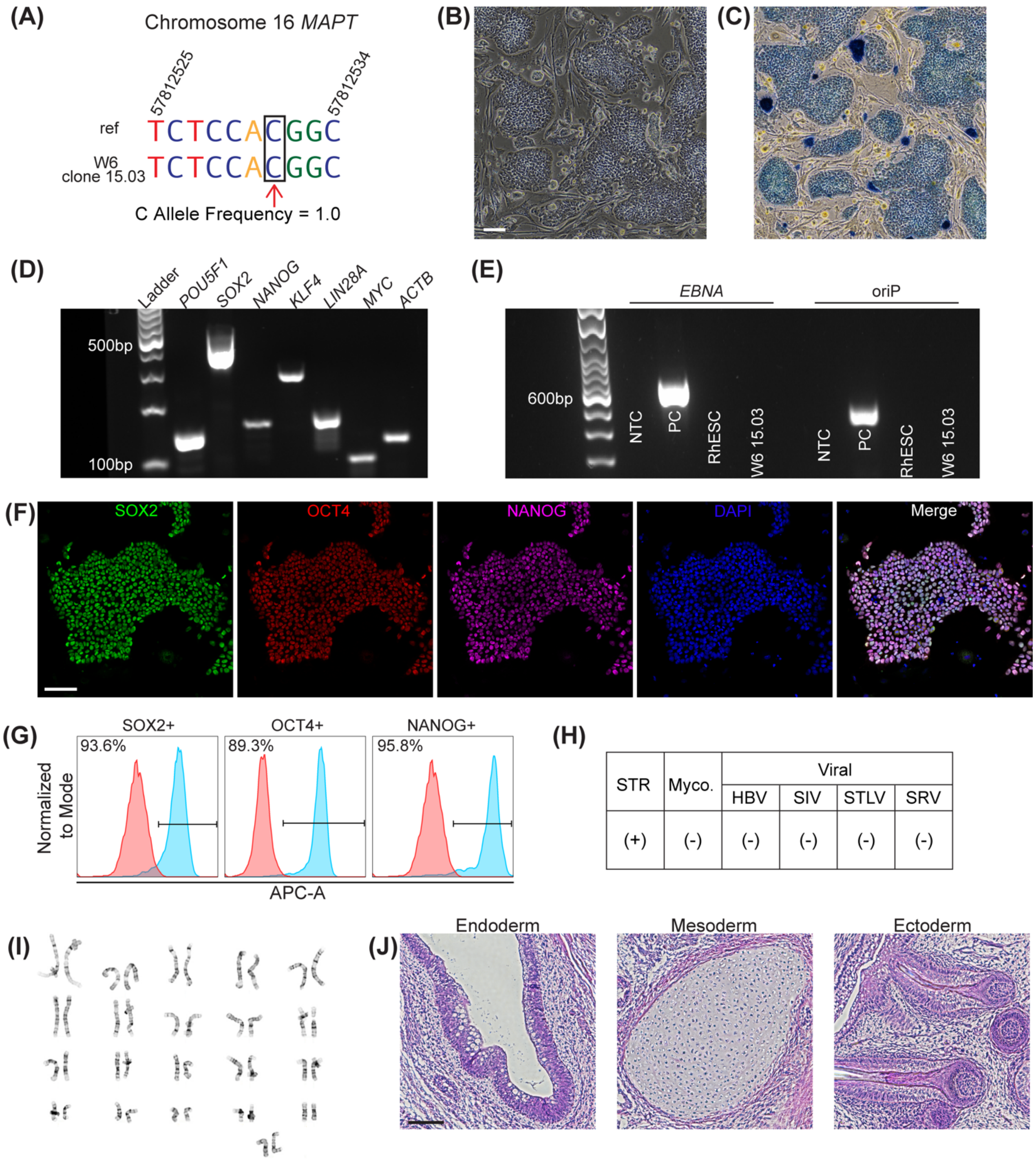


**Supplementary Figure 4.** Characterization of RhiPSC line W6 clone 15.03. (A) Amplicon sequencing of *MAPT* R406W with wild-type C allele indicated in the black box with a red arrow compared to reference genome (ref). (B) Phase contrast image of colonies grown on MEFs and (C) positive alkaline phosphatase staining; scale bar, 100 µm. (D) RT-PCR detection of endogenous pluripotent mRNAs *POU5F1*, *SOX2*, *NANOG*, *KLF4*, *LIN28*, and *MYC*. *ACTB* was used as a reference gene. (E) Amplification of EBNA and oriP in RhiPSCs at P11, no template control (NTC), and the rhesus embryonic stem cell line r420 (RhESC). Plasmid EM2K was used as a positive control (PC) for the detection of EBNA and oriP. (F) Positive immunofluorescence staining for pluripotency proteins SOX2 (green), OCT4 (red), and NANOG (far red), colocalized with DAPI (blue). Scale bar, 100 µm. (G) Flow cytometry-detected APC-A+ cells were 93.6% positive for SOX2, 89.3% positive for OCT4, and 95.8% positive for NANOG. (H) Short tandem repeat (STR) analysis confirmed (+) RhiPSC line identity. Mycoplasma (Myco.), Herpes B Virus (HBV), Simian Immunodeficiency Virus (SIV), Simian T-Lymphotropic Virus (STLV), and Simian Retrovirus (SRV) were not detected (-). (I) Normal 42,XX karyotype at P13. (J) H&E staining of a teratoma containing endoderm, mesoderm, and ectoderm. Scale bar, 100 µm.

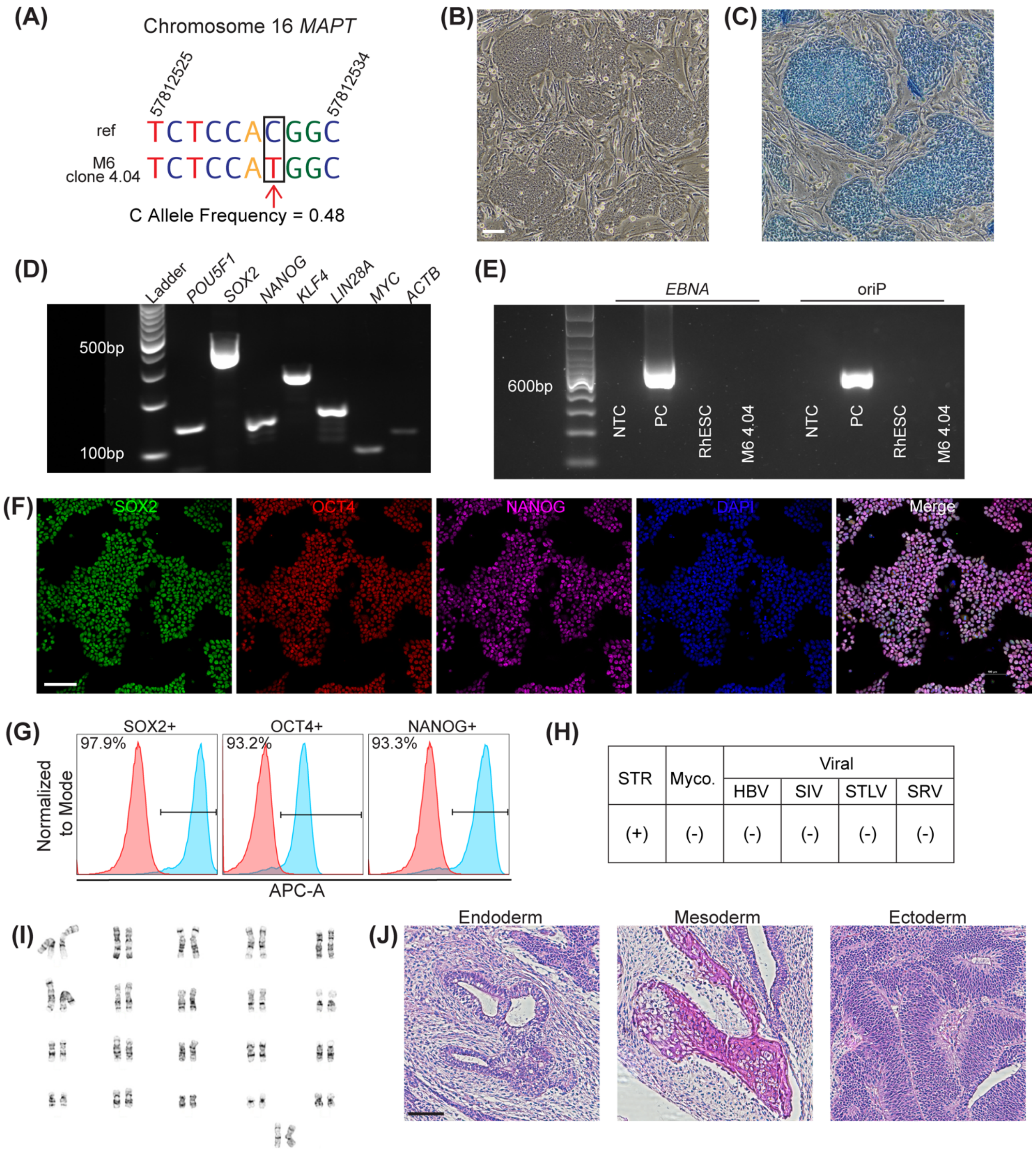


**Supplementary Figure 5.** Characterization of RhiPSC line M6 clone 4.04. (A) Amplicon sequencing of *MAPT* R406W with mutant T allele indicated in the black box with a red arrow compared to reference genome (ref). (B) Phase contrast image of colonies grown on MEFs and (C) positive alkaline phosphatase staining; scale bar, 100 µm. (D) RT-PCR detection of endogenous pluripotent mRNAs *POU5F1*, *SOX2*, *NANOG*, *KLF4*, *LIN28*, and *MYC*. *ACTB* was used as a reference gene. (E) Amplification of EBNA and oriP in RhiPSCs at P14, no template control (NTC), and the rhesus embryonic stem cell line r420 (RhESC). Plasmid EM2K was used as a positive control (PC) for the detection of EBNA and oriP. (F) Positive immunofluorescence staining for pluripotency proteins SOX2 (green), OCT4 (red), and NANOG (far red), colocalized with DAPI (blue). Scale bar, 100 µm. (G) Flow cytometry-detected APC-A+ cells were 97.9% positive for SOX2, 93.2% positive for OCT4, and 93.3% positive for NANOG. (H) Short tandem repeat (STR) analysis confirmed (+) RhiPSC line identity. Mycoplasma (Myco.), Herpes B Virus (HBV), Simian Immunodeficiency Virus (SIV), Simian T-Lymphotropic Virus (STLV), and Simian Retrovirus (SRV) were not detected (-). (I) Normal 42,XX karyotype at P13. (J) H&E staining of a teratoma containing endoderm, mesoderm, and ectoderm. Scale bar, 100 µm.

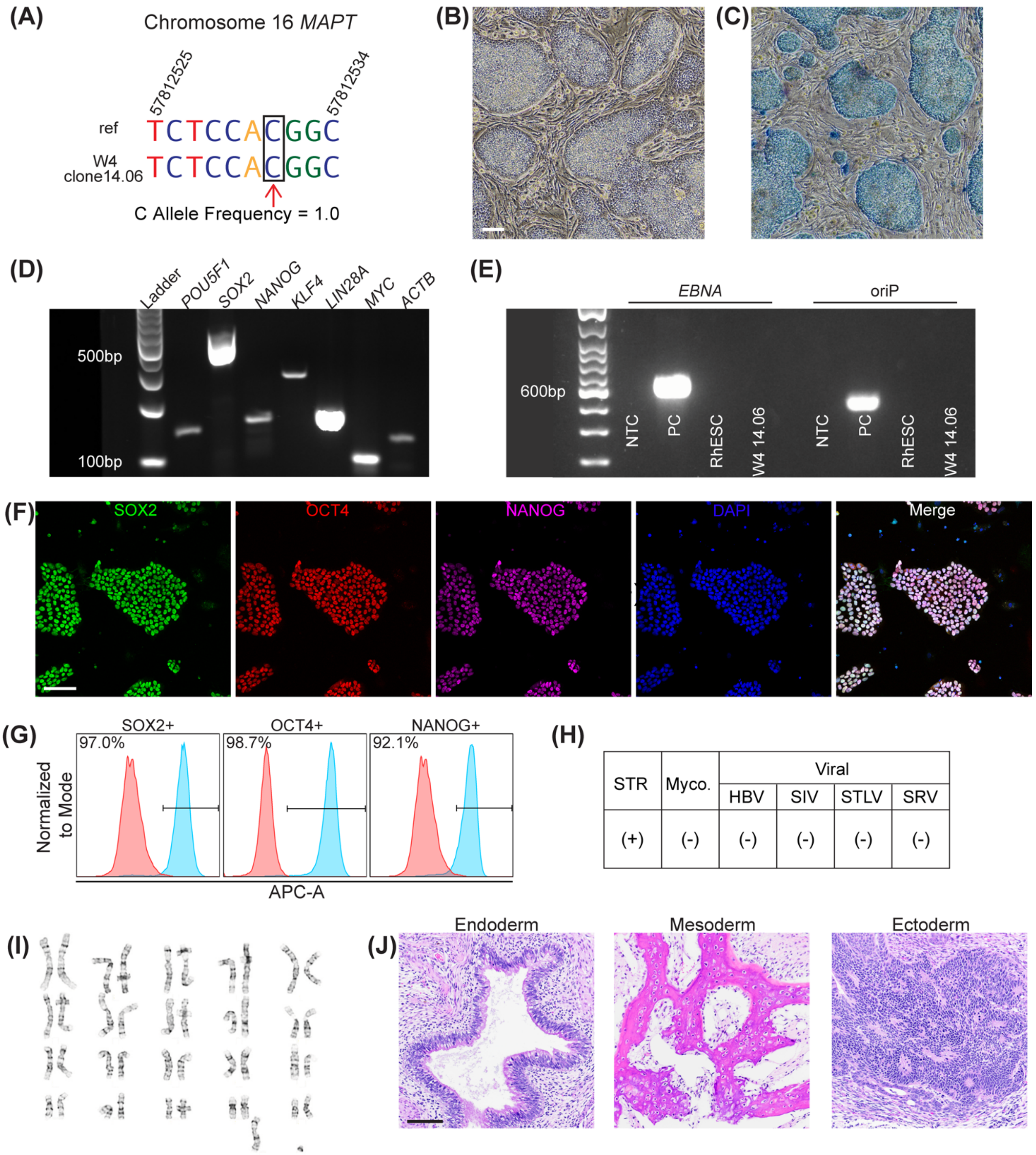


**Supplementary Figure 6.** Characterization of RhiPSC line W4 clone 14.06. (A) Amplicon sequencing of *MAPT* R406W with wild type C allele indicated in the black box with a red arrow compared to reference genome (ref). (B) Phase contrast image of colonies grown on MEFs and (C) positive alkaline phosphatase staining; scale bar, 100 µm. (D) RT-PCR detection of endogenous pluripotent mRNAs *POU5F1*, *SOX2*, *NANOG*, *KLF4*, *LIN28*, and *MYC*. *ACTB* was used as a reference gene. (E) Amplification of EBNA and oriP in RhiPSCs at P14, no template control (NTC), and the rhesus embryonic stem cell line r420 (RhESC). Plasmid EM2K was used as a positive control (PC) for the detection of EBNA and oriP. (F) Positive immunofluorescence staining for pluripotency proteins SOX2 (green), OCT4 (red), and NANOG (far red), colocalized with DAPI (blue). Scale bar, 100 µm. (G) Flow cytometry-detected APC-A+ cells were 97% positive for SOX2, 98.7% positive for OCT4, and 92.1% positive for NANOG. (H) Short tandem repeat (STR) analysis confirmed (+) RhiPSC line identity. Mycoplasma (Myco.), Herpes B Virus (HBV), Simian Immunodeficiency Virus (SIV), Simian T-Lymphotropic Virus (STLV), and Simian Retrovirus (SRV) were not detected (-). (I) Normal 42,XY karyotype at P10. (J) H&E staining of a teratoma containing endoderm, mesoderm, and ectoderm. Scale bar, 100 µm.

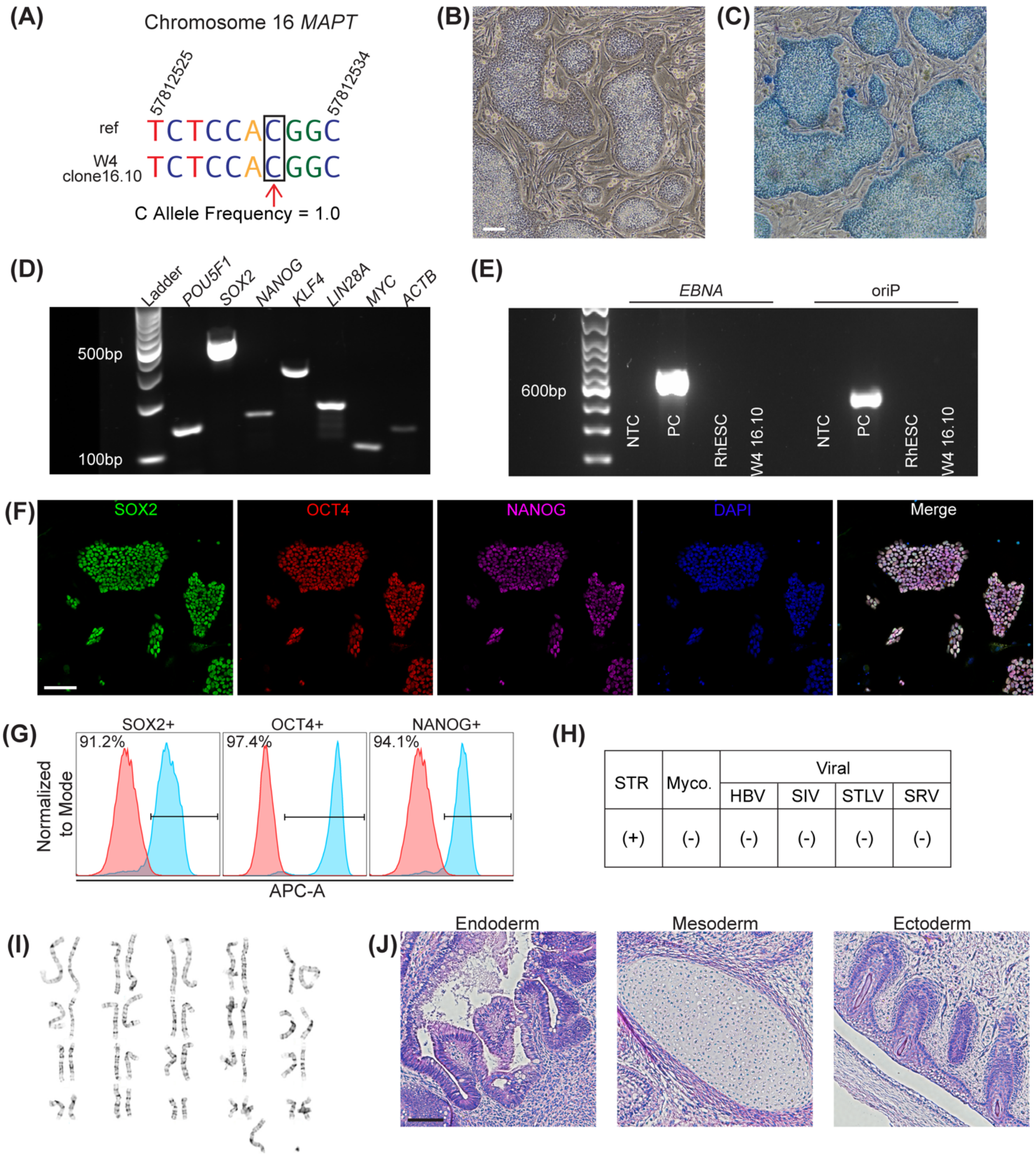


| STR | Myco. | Viral | | | |
|---|---|---|---|---|---|
| | | HBV | SIV | STLV | SRV |
| (+) | (-) | (-) | (-) | (-) | (-) |

**Supplementary Figure 7**. Characterization of RhiPSC line W4 clone 16.10. (A) Amplicon sequencing of *MAPT* R406W with wild type C allele indicated in the black box with a red arrow compared to reference genome (ref). (B) Phase contrast image of colonies grown on MEFs and (C) positive alkaline phosphatase staining; scale bar, 100 µm. (D) RT-PCR detection of endogenous pluripotent mRNAs *POU5F1*, *SOX2*, *NANOG*, *KLF4*, *LIN28*, and *MYC*. *ACTB* was used as a reference gene. (E) Amplification of EBNA and oriP in RhiPSCs at P14, no template control (NTC), and the rhesus embryonic stem cell line r420 (RhESC). Plasmid EM2K was used as a positive control (PC) for the detection of EBNA and oriP. (F) Positive immunofluorescence staining for pluripotency proteins SOX2 (green), OCT4 (red), and NANOG (far red), colocalized with DAPI (blue). Scale bar, 100 µm. (G) Flow cytometry-detected APC-A+ cells were 91.2% positive for SOX2, 97.4% positive for OCT4, and 94.1% positive for NANOG. (H) Short tandem repeat (STR) analysis confirmed (+) RhiPSC line identity. Mycoplasma (Myco.), Herpes B Virus (HBV), Simian Immunodeficiency Virus (SIV), Simian T-Lymphotropic Virus (STLV), and Simian Retrovirus (SRV) were not detected (-). (I) Normal 42,XY karyotype at P10. (J) H&E staining of a teratoma containing endoderm, mesoderm, and ectoderm. Scale bar, 100 µm.

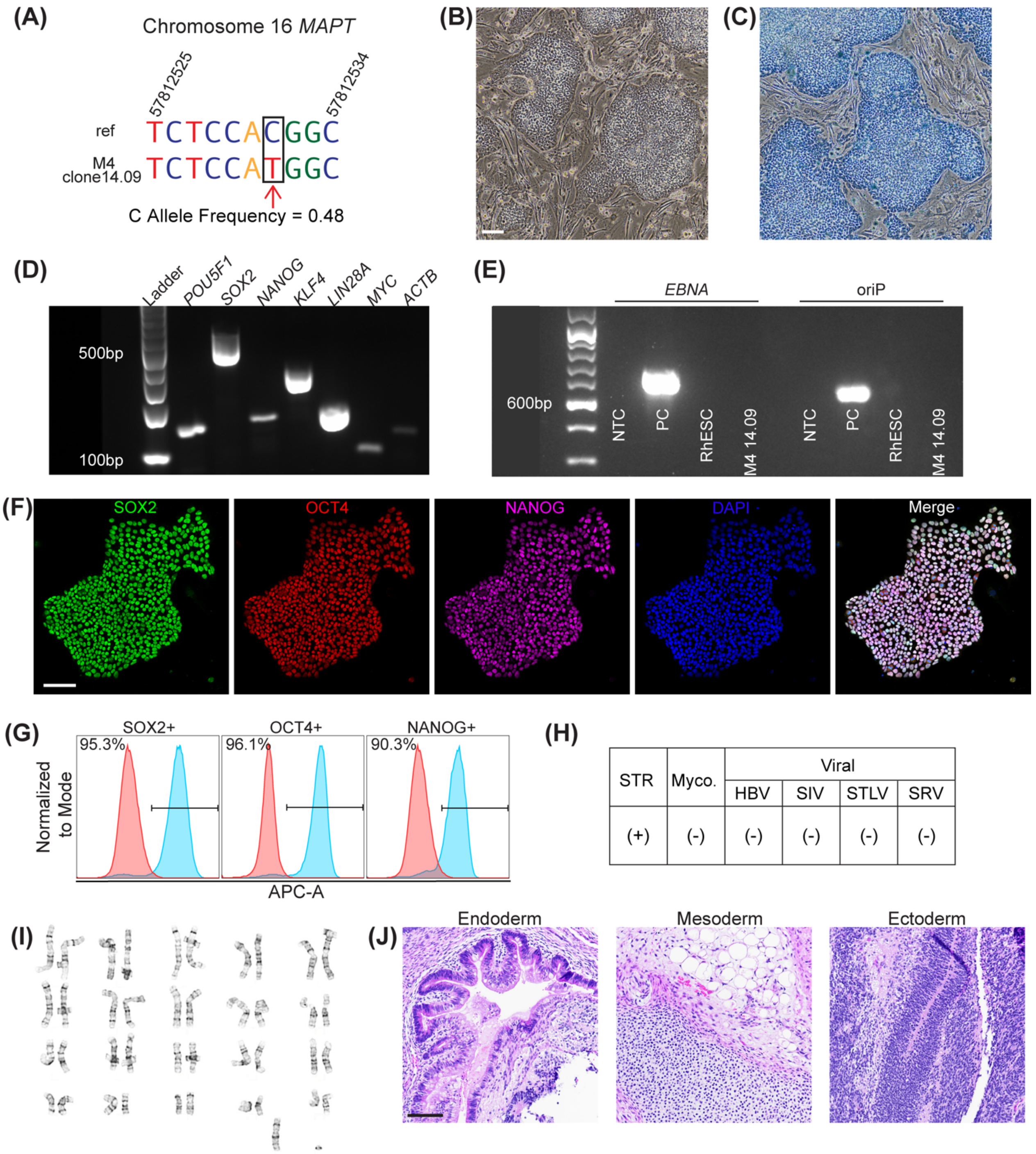


**Supplementary Figure 8.** Characterization of RhiPSC line M4 clone 14.09. (A) Amplicon sequencing of *MAPT* R406W with mutant T allele indicated in the black box with a red arrow compared to reference genome (ref). (B) Phase contrast image of colonies grown on MEFs and (C) positive alkaline phosphatase staining; scale bar, 100 µm. (D) RT-PCR detection of endogenous pluripotent mRNAs *POU5F1*, *SOX2*, *NANOG*, *KLF4*, *LIN28*, and *MYC*. *ACTB* was used as a reference gene. (E) Amplification of EBNA and oriP in RhiPSCs at P14, no template control (NTC), and the rhesus embryonic stem cell line r420 (RhESC). Plasmid EM2K was used as a positive control (PC) for the detection of EBNA and oriP. (F) Positive immunofluorescence staining for pluripotency proteins SOX2 (green), OCT4 (red), and NANOG (far red), colocalized with DAPI (blue). Scale bar, 100 µm. (G) Flow cytometry-detected APC-A+ cells were 95.3% positive for SOX2, 96.1% positive for OCT4, and 90.3% positive for NANOG. (H) Short tandem repeat (STR) analysis confirmed (+) RhiPSC line identity. Mycoplasma (Myco.), Herpes B Virus (HBV), Simian Immunodeficiency Virus (SIV), Simian T-Lymphotropic Virus (STLV), and Simian Retrovirus (SRV) were not detected (-). (I) Normal 42,XY karyotype at P18. (J) H&E staining of a teratoma containing endoderm, mesoderm, and ectoderm. Scale bar, 100 µm.

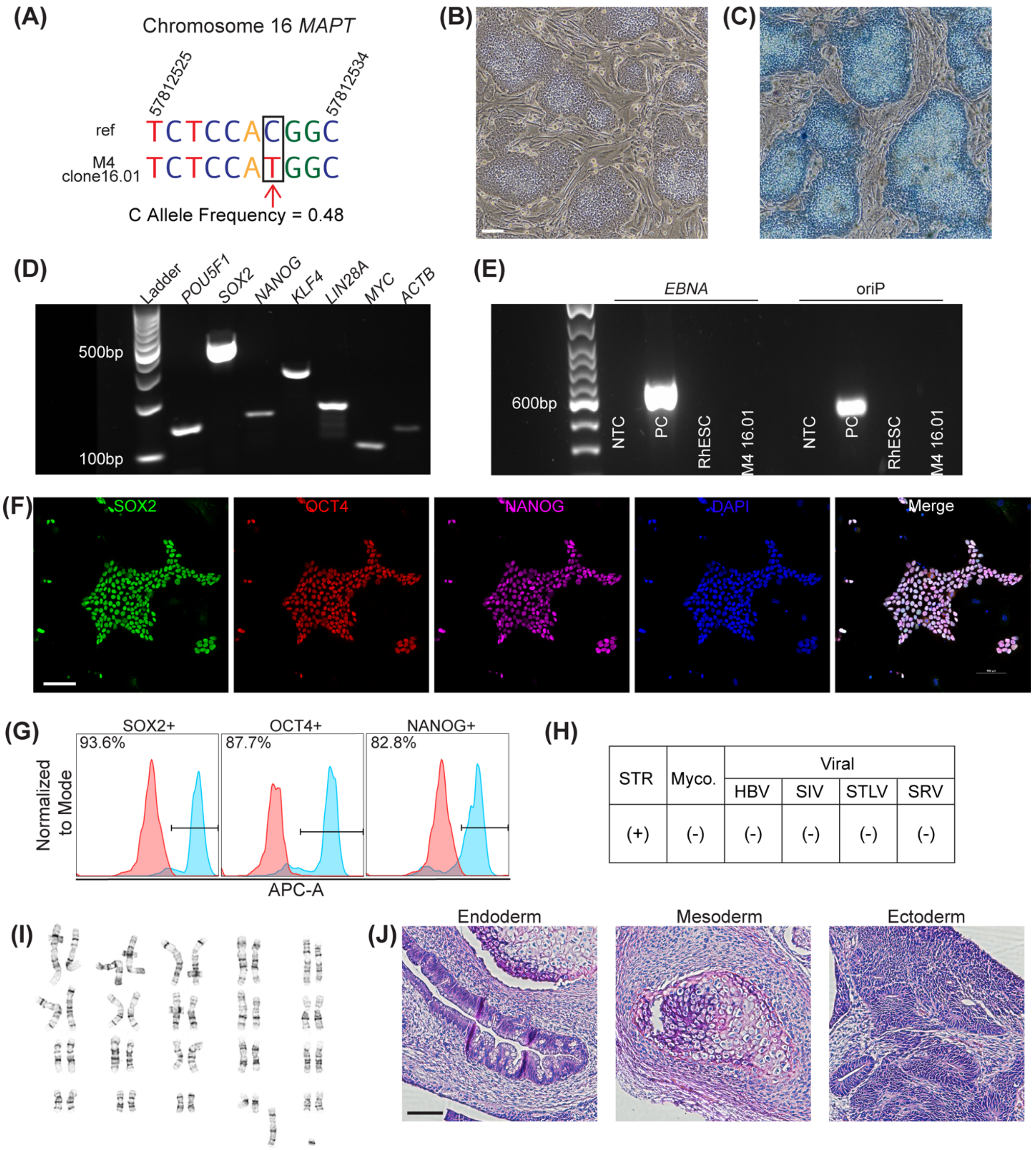


**Supplementary Figure 9**. Characterization of RhiPSC line M4 clone 16.01. (A) Amplicon sequencing of *MAPT* R406W with mutant T allele indicated in the black box with a red arrow compared to reference genome (ref). (B) Phase contrast image of colonies grown on MEFs and (C) positive alkaline phosphatase staining; scale bar, 100 µm. (D) RT-PCR detection of endogenous pluripotent mRNAs *POU5F1*, *SOX2*, *NANOG*, *KLF4*, *LIN28*, and *MYC*. *ACTB* was used as a reference gene. (E) Amplification of EBNA and oriP in RhiPSCs at P14, no template control (NTC), and the rhesus embryonic stem cell line r420 (RhESC). Plasmid EM2K was used as a positive control (PC) for the detection of EBNA and oriP. (F) Positive immunofluorescence staining for pluripotency proteins SOX2 (green), OCT4 (red), and NANOG (far red), colocalized with DAPI (blue). Scale bar, 100 µm.(G) Flow cytometry-detected APC-A+ cells were 93.6% positive for SOX2, 87.7% positive for OCT4, and 82.8% positive for NANOG. (H) Short tandem repeat (STR) analysis confirmed (+) RhiPSC line identity. Mycoplasma (Myco.), Herpes B Virus (HBV), Simian Immunodeficiency Virus (SIV), Simian T-Lymphotropic Virus (STLV), and Simian Retrovirus (SRV) were not detected (-). (I) Normal 42,XY karyotype at P18. (J) H&E staining of a teratoma containing endoderm, mesoderm, and ectoderm. Scale bar, 100 µm.
